# Base Pairing and Functional Insights into *N^3^*-methylcytidine (m^3^C) in RNA

**DOI:** 10.1101/2020.08.27.270322

**Authors:** Song Mao, Phensinee Haruehanroengra, Srivathsan V. Ranganathan, Fusheng Shen, Thomas J. Begley, Jia Sheng

## Abstract

*N^3^*-methylcytidine (m^3^C) is present in both eukaryotic tRNA and mRNA and plays critical roles in many biological processes. We report the synthesis of the m^3^C phosphoramidite building block and its containing RNA oligonucleotides. The base-pairing stability and specificity studies show that the m^3^C modification significantly disrupts the stability of the Watson-Crick C:G pair. Further m^3^C decreases the base pairing discrimination between C:G and the other mismatched C:A, C:U, and C:C pairs. Our molecular dynamic simulation study further reveals the detailed structural insights into the m^3^C:G base pairing pattern in an RNA duplex. More importantly, the biochemical investigation of m^3^C using reverse transcription shows that *N^3^*-methylation specifies the C:A pair and induces a G to A mutation using HIV-1-RT, MMLV-RT and MutiScribe™-RT enzymes, all with relatively low replication fidelity. For other reverse transcriptases with higher fidelity like AMV-RT, the methylation could completely shut down DNA synthesis.

## INTRODUCTION

Natural RNA systems in all organism, from the simplest prokaryote *Nanoarchaeum equitans* to humans, utilize the four regular nucleosides (adenosine, guanosine, cytidine, and uridine) and a variety of post-transcriptional modifications to achieve structural and functional specificity and diversity. (Nachtergaele and He, 2017) To date, more than 160 distinct chemical modifications that decorate different positions of nucleobases, ribose and phosphate backbone in RNA nucleotides have been discovered, (Basanta-Sanchez et al., 2016; Boccaletto et al., 2018; Cantara et al., 2011; Machnicka et al., 2013; Wu et al., 2020) since the first discovery of the modified nucleoside in the 1950s. (Holley et al., 1965a; Holley et al., 1965b) Many RNA modifications have been demonstrated to play critical roles in both normal and disease cellular functions and processes, such as development, circadian rhythms, embryonic stem cell differentiation, meiotic progression, temperature adaptation, stress response and tumorigenesis, etc. (Machnicka et al., 2013) Similar to DNA and protein based epigenetic markers, RNA modifications (also termed the ‘epitranscriptome’) are dynamically and reversibly regulated by specific reader, writer and eraser enzymes, representing a new layer of gene regulation. (Roundtree et al., 2017) Accordingly, RNA modification-associated enzymes, as an important research frontier towards RNA-based drug discovery, have become useful molecular tools and drug targets. (Jiang et al., 2017)

Benefiting from a variety of recently developed chemical biology tools and high-throughput detection strategies, RNA methylation has been identified in different RNAs from all organisms.(Clark et al., 2016; Hori, 2014; Mongan et al., 2019; Nachtergaele and He, 2018; Roundtree et al., 2017; Sergiev et al., 2018; Song and Yi, 2017) In addition, corresponding writers and erasers and bonding proteins (“readers’) (Shi et al., 2019) have been identified for many RNA methylation and been shown to impact numerous biological functions and diseases processes. For example, the tRNA methylations 5-methylcytidine (m^5^C), *N^1^*-methylguanidine (m^1^G), *N^1^*-methyladenosine (m^1^A), *N^7^*-methylguanidine (m^7^G) and 2’-O-methylated sugar (2’-Nm) in the anticodon stem loops of transfer RNA (tRNA) are directly involved in the codon recognition and can induce or inhibit frameshifting mutations during translation. (Fu et al., 2014; Wang et al., 2014) In addition, *N^6^*-methyladenosine (m^6^A), the most abundant internal mRNA methylation, is linked to numerous biological functions, including mRNA stability, RNA structure switches, mRNA splicing, RNA export, translation and miRNA biogenesis. (Desrosiers et al., 1974; Song and Yi, 2017; Zaccara et al., 2019) Moreover, RNA methylation also has been found in viral RNA, which impacts viral gene expression and has great potential for stimulating therapeutic developments. (Chen et al., 2019a; Ciuffi, 2016; Lichinchi et al., 2016; Wu, 2019)

*N^3^*-methylcytidine (m^3^C), first discovered in *Saccharomyces cerevisiae* total RNA (Hall, 1963) and later found in eukaryotic tRNA (Clark et al., 2016; Cozen et al., 2015; D’Silva et al., 2011; Han et al., 2017; Iwanami and Brown, 1968; Noma et al., 2011; Olson et al., 1981), occurs most frequently in anticodon stem loops to impact tRNA structure, affinity for the ribosome and decoding activity. The working enzymes responsible for m^3^C in tRNA are the methyltransferase Trm140 or the complex of Trm140 and Trm141 (METTL2 and METTL6). Recently, Fu and coworkers reported the new discovery of METTL8 as an mRNA m^3^C writer enzyme and provided the first evidence of the existence of m^3^C modification in the mRNA of mice and humans, (Liu and He, 2017; Xu et al., 2017) The m^3^C modification may play versatile roles in impacting mRNA processing and biological functions. More interestingly, m^3^C has been uniquely detected in the viral RNAs from Huh7, ZIKV and DENV virions and the cells with these virus infections. (McIntyre et al., 2018) The m^3^C modification also has the potential to be demethylated by eraser enzymes. Alkylation repair homolog 3 (ALKBH3), as well as its bacterial ancestor Alkb, have been shown to demethylate the m^3^C in tRNA to affect RNA stability and prevent degradation.(Chen et al., 2019b; Ougland et al., 2004) ALKBH3 expression has also been linked to tumor progression and the regulation of protein synthesis, suggesting m^3^C plays a prominent role in cancer biology. (Ueda et al., 2017)

Although much effort went in to the discovery and detection of m^3^C, little is known about its fundamental properties and biological functions. Since the *N^3^*-position directly participates in the Watson-Crick pairing, this methylation is expected to disrupt the C:G pair and reduce the base pairing fidelity of cytosine. In addition, the methyl group on m^3^C might also regulate binding by RNA readers. Therefore, we hypothesize that the methylation at *N^3^*-position of cytidine is a cellular mechanism to modulate base pairing specificity and affect the efficiency and fidelity of transcription and reverse-transcription, thus increasing mutation rates, which could be beneficial to certain biological systems like virus. To the best of our knowledge, no chemical synthesis and base-pairing studies of RNA oligonucleotides containing m^3^C modification have been reported. In this work, we report the new chemical synthesis of m^3^C phosphoramidite building block and its incorporation into RNA oligonucleotides. The subsequent base-pairing stability and specificity studies of RNA duplexes containing one and two m^3^C residues at different positions support the idea that the m^3^C decreases both duplex stability and base pairing discrimination between C:G pair and other mismatched pairs. Our molecular dynamic simulation study further provides detailed structural insights into the m^3^C:G base pairing pattern in an RNA duplex. Furthermore, we used m^3^C in reverse transcription assays in the presence of AMV-RT, HIV-1-RT, MMLV-RT and MutiScribe™-RT and found that this methylation could specify the C:A pair for some RT enzymes with low fidelity, which would induce the G to A mutation. For reverse transcriptase enzymes with higher fidelity (i.e., AMV-RT), m^3^C could completely shut down DNA synthesis.

## RESULTS AND DISCUSSION

### Chemical synthesis of m^3^C phosphoramidite building block and its containing RNA oligonucleotides

Although the synthesis of m^3^C nucleoside has been achieved, (Brookes and Lawley, 1962; Ogilvie and Kader, 1983) more general phosphoramidite building blocks are required to make different scales of RNA strands through the solid phase synthesis of oligonucleotides. We started the synthesis of m^3^C from the commercially available cytidine (**1**, **scheme 1**), which was directly methylated by using MeI without any base to obtain the m^3^C nucleoside. The sequential protections of the 5’-hydroxyl with dimethoxyltrityl (DMTr) group and *N^4^* position with benzoyl (Bz) group yielded compound **4**. Subsequently, the 2’-hydroxyl group was protected with *tert*-butyldimethylsilyl (TBDMS) group to obtain compound **5**, which is the key intermediate to make the final phosphoramidite building block **6** for the oligonucleotides solid phase synthesis.

**Scheme 1.**
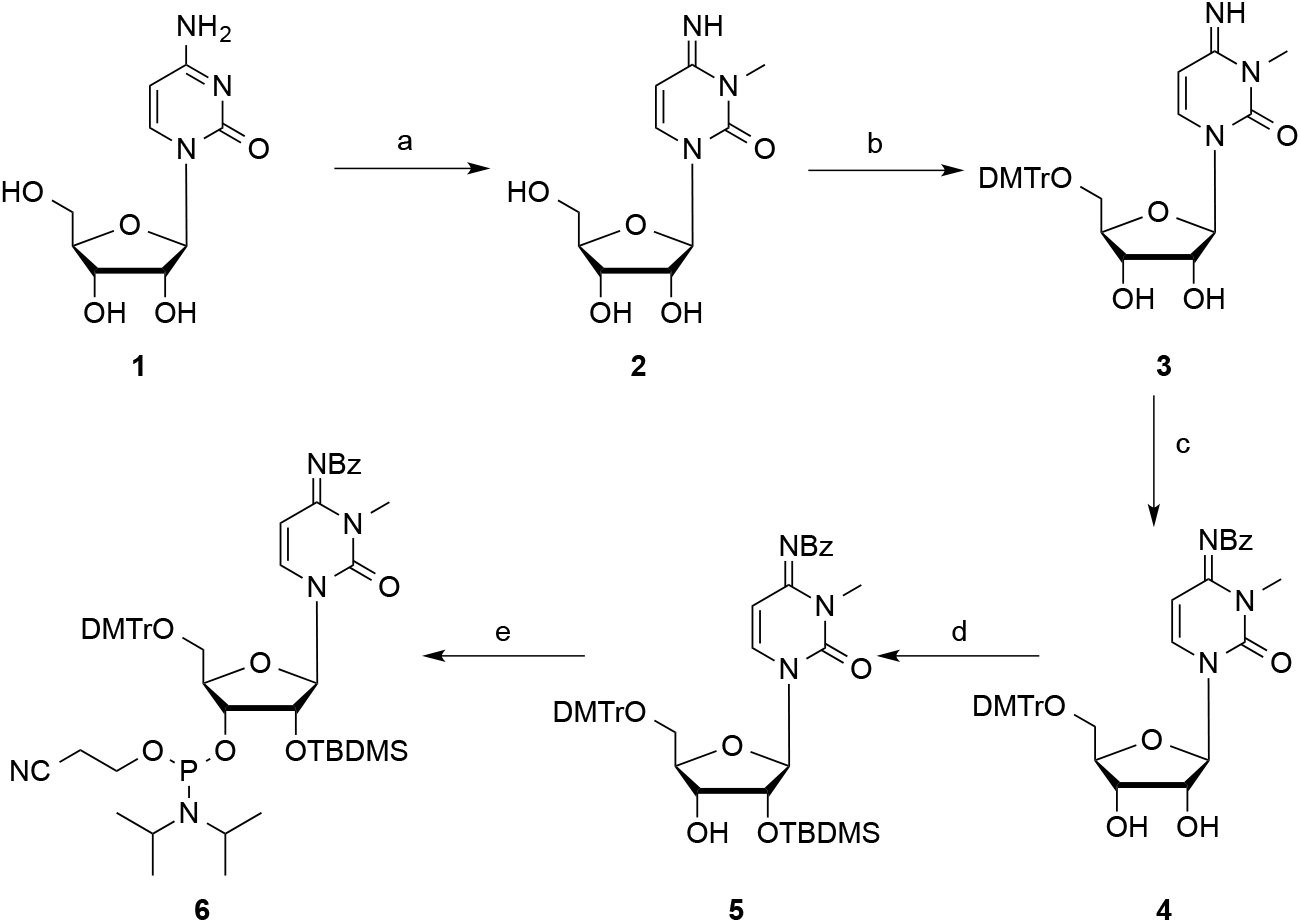
Synthesis of *N^3^*-methyl-cytidine phosphoramidite **10**. Reagents and conditions: (a) MeI, DMF; (b) DMTrCl, Py; (c) TMSCl, Py; BzCl; (d) TBDMSCl, imidazole, DMF; (e) (*i*-Pr_2_N)_2_P(Cl)OCH_2_CH_2_CN, (*i*-Pr)_2_NEt, 1-methylimidazole, DCM.

As expected, the m^3^C phosphoramidite building block is well compatible with the solid phase synthesis conditions including the trichloroacetic acid (TCA) and oxidative iodine treatments, resulting in very similar coupling yields as the commercially available native phosphoramidites. They are also stable in the basic cleavage from the solid phase beads and the Et_3_N•3HF treatment to remove TBDMS protecting groups during the RNA oligonucleotide deprotection and purification. As the demonstration, six RNA strands containing this modification have been synthesized and confirmed by ESI- or MALDI-MS, as shown in **Table 1**.

**Table 1.** RNA sequences containing m^3^C.

| Entry | RNA Sequences | Measured (calc.) m/z |
| --- | --- | --- |
| ON1 | 5'-AAUGCm<sup>3</sup>CGCACUG-3' | [M+H]<sup>+</sup> = 3807.5 (3807.6) |
| ON2 | 5'-GGACUm<sup>3</sup>CCUGCAG-3' | [M+H]<sup>+</sup> = 3823.6 (3823.6) |
| ON3 | 5'-Um<sup>3</sup>CGUACGA-3' | [M+H]<sup>+</sup> = 2523.1 (2522.4) |
| ON4 | 5'-GUAm<sup>3</sup>CGUAC-3' | [M+H]<sup>+</sup> = 2522.5 (2522.4) |
| ON5 | 5'-CCGGm<sup>3</sup>CGCCGG-3' | [M+H]<sup>+</sup> = 3203.7 (3203.5) |
| ON6 | 5'-CGCGAAUUm<sup>3</sup>CGCG-3' | [M+H]<sup>+</sup> = 3823.6 (3823.6) |

### Thermal denaturation and base pairing studies of m^3^C RNA duplexes

We synthesized two sets of RNA strands to investigate the thermodynamic properties and base pairing specificity of m^3^C containing RNA duplexes. The normalized *T*_m_ curves of native and modified RNA duplexes, [5’-GGACU<u>X</u>CUGCAG-3’ & 3’-CCUGA<u>Y</u>GACGUC-5’] with Watson-Crick and other non-canonical base pairs (X pairs with Y), are shown in **Figure 1**. The detailed melting temperature data are summarized in **Table 2**. Compared to the native counterparts, m^3^C-modified RNA duplexes showed dramatically decreased thermal stability. In the native C:G paired 12-mer duplexes (compare entry 2 and 7), the m^3^C decreases the *T*_m_ by 19.7 °C, corresponding to a ΔG^0^ reduction of 9.6 kcal/mol. Similarly, the non-canonical base paired (ex. C:A, C:U and C:C) duplexes containing this modification also showed significantly lower melting temperatures. The *T*_m_ drops by 9.9 °C in the C:A mismatched duplex (entry 3 *vs* 8), 7.0 °C in the C:U mismatched one (entry 4 *vs* 9) and 4.0 °C for the C:C mismatched one (entry 5 *vs* 10), corresponding to the ΔG^0^ reduction of 4.5, 4.2 and 2.5 kcal/mol respectively.

**Figure 1.**
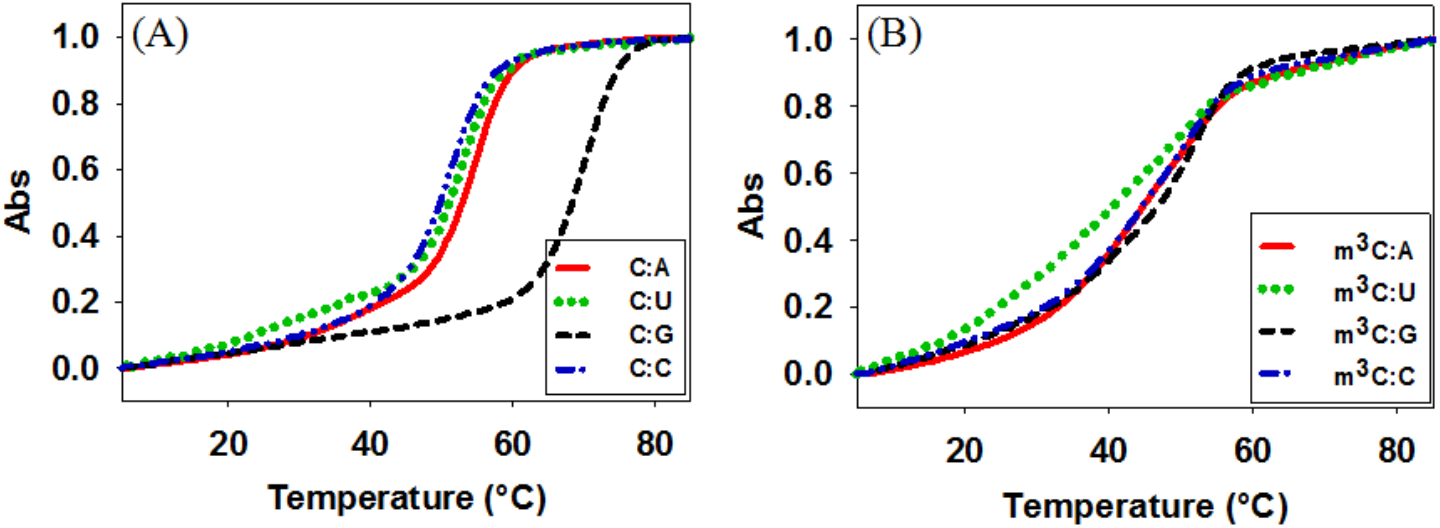
Normalized UV-melting curves of RNA duplexes. (A) Native sequence 5’-GGACU<u>C</u>CUGCAG-3’) pairs with matched and mismatched strands. (B) m^3^C modification sequence (5’-GGACU<u>m^3^C</u>CUGCAG-3’) pairs with matched and mismatched sequences.

These results indicate that m^3^C modification significantly disrupts the C:G pair and the overall duplex stability. Indeed, when we compared this modification with the native C in a self-complementary 10-mer duplex context (CCGG<u>C*</u>GCCGG)_2_, where two consecutive m^3^C:G are introduced in the middle of the duplex, the *T*_m_ drops by 35.7 °C (entry 11 *vs* 12, **Table 2**), as shown in **Figure S34**. On the other hand, the comparison of base pairing specificity in this duplex system indicated that m^3^C decreases the discrimination between C:G pair and other mismatched C:A, C:U and C:C pairs (entries 7-9, **Table 2**). The lowest *T*_m_ difference is 3.2 °C between m^3^C:G-duplex and m^3^C:C-one, and the highest *T*_m_ difference is only 5.6 °C between m^3^C:G-duplex and m^3^C:A-one. In comparison, in the nonmodified native RNA duplexes, these *T*_m_ differences vary from 15.4 to 18.9 °C (entries 2-5), dramatically bigger than the modified m^3^C counterparts.

**Table 2.**
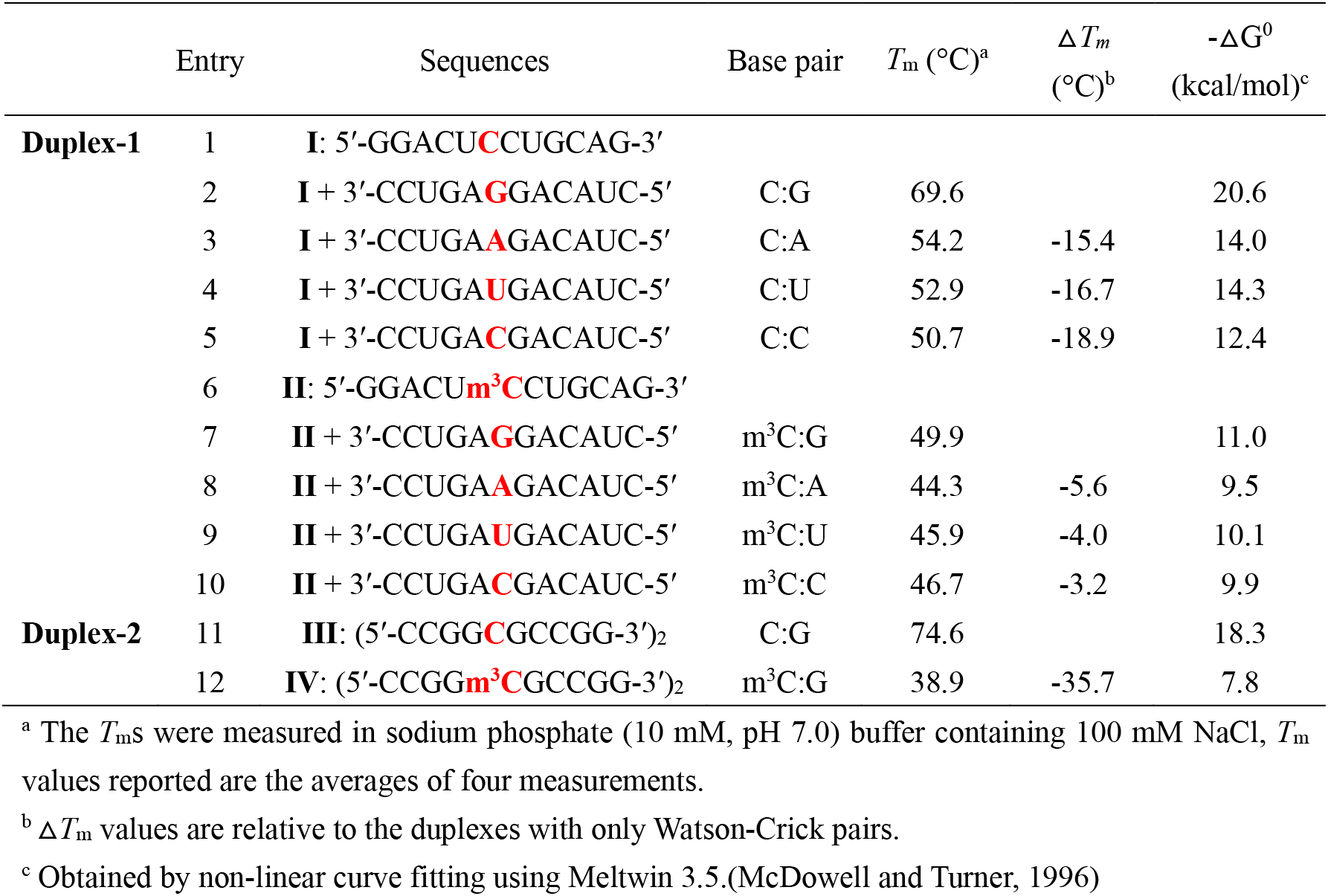
Melting temperatures of native and m^3^C-modified RNA duplexes.

In addition, we also evaluated the impact of multiple m^3^C modifications at different positions using a longer 22-mer RNA duplex template. As shown in **Table 3** and **Figure S35**, the *T*_m_ of RNA duplex containing a single m^3^C residue at C17 position (close to the 3’ end) drops by 5.5 °C (entry 2 *vs* 4, **Table 3**). In comparison, the *T*_m_ of the one containing two m^3^C residues at C17 and C19 positions decreases by only 6.5 °C (entry 2 *vs* 4 *vs* 6, **Table 3**), meaning that the additional m^3^C residue in the adjacent position has a small impact on duplex stability with only a 1.0 °C of *T*_m_ decrease and the further structural perturbation is ‘buffered’ by the first m^3^C modification. By contrast, separating the two m^3^C modifications at C5 and C17 positions resulted in dramatic drop of *T*_m_ by 37.2 °C (entry 2 *vs* 8, **Table 3**). While it has been known that RNA duplex structure is flexible to accommodate many different chemical modifications, the wide duplex stability range that m^3^C could induce and the capability of fine-tuning *T*_m_ in a position-dependent manner might be useful for therapeutic applications, to enhance the efficacy of antisense and RNAi based mechanisms.

**Table 3.**
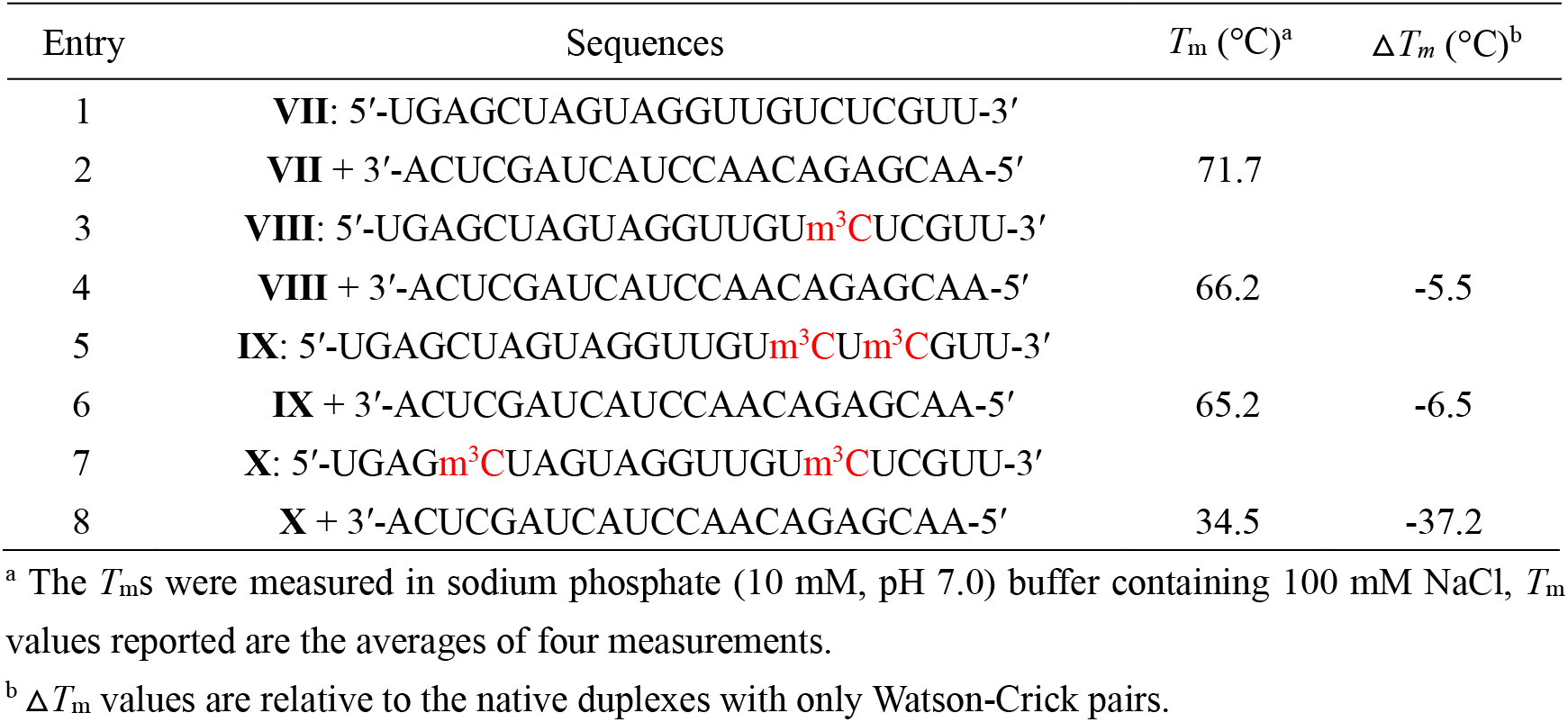
Melting temperatures of native and m^3^C-modified 22-mer RNA duplexes.

### Molecular simulation studies of m^3^C-RNA duplex

In order to investigate more detailed structural insights into the base pairing patterns of the m^3^C containing RNA duplex, we conducted molecular dynamic simulation studies. The results from the MD simulations are summarized in **Figure 2**. The RNA duplex was simulated in both canonical (C-G) and modified (m^3^C-G) forms. We calculated the hydrogen-bonding distances between the donor-acceptor pairs for the canonical and modified base-pairs from the ensemble of structures generated in the production run. The curves are moved vertically for visual clarity. The canonical C:G pair retains all the three hydrogen bonds throughout the simulation. However, in the modified m^3^C-G pair, the m^3^C rotates for about 45 degrees to fully expose the methyl group into the major groove and avoid the clashing with N^1^ of the pairing G. The conformational change observed in the m^3^C-G pair also allows for the single hydrogen bond acceptor (O^2^ of m^3^C) to form bifurcated hydrogen bonds with N^1^ and N^2^ of guanine. The bifurcated hydrogen bonds are weaker than the normal one as evidenced by the higher average distances and fluctuations. Similar bonding patterns might also exist in other mis-matched pairs to minimize the discrimination of C to other bases. Overall, we observed weakening of hydrogen bonding in the modified m^3^C-G base-pair, which is consistent with our thermostability studies.

**Figure 2.**
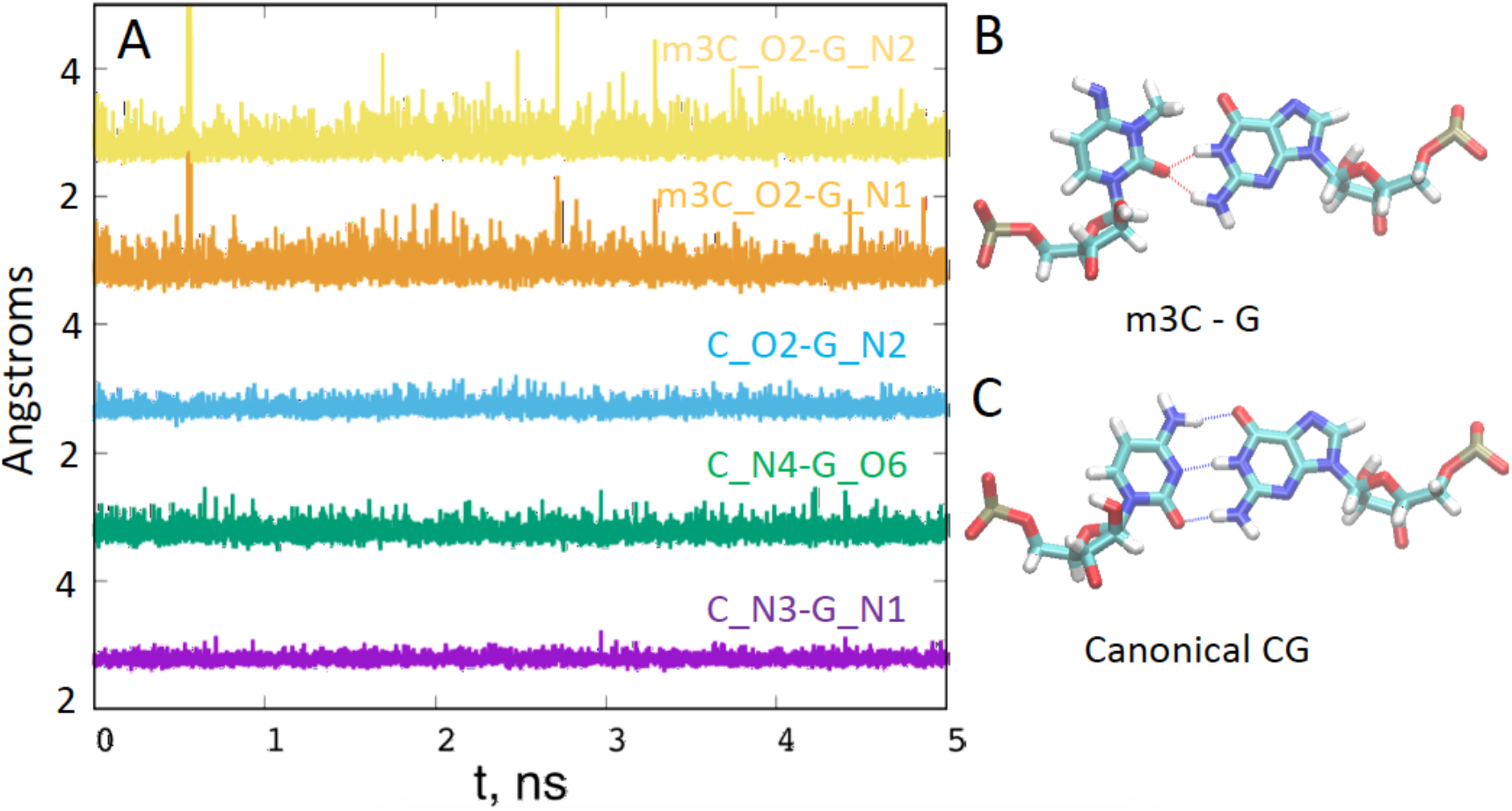
Molecular simulation results. The hydrogen bonding distances (A) vary at different time points in the RNA duplex containing m^3^C:G (B) and native C:G (C) pairs.

### Impacts of m^3^C modification on reverse transcription

In order to study the potential biological consequences of m^3^C induced changes in hydrogen bonding during nucleic acid synthesis, we conducted reverse transcription assays using a modified-template directed primer extension reaction, as shown in **Figure 3**. The 5’-end of DNA primer was labeled with the fluorescent FAM group and a 31 nt-long modified RNA was used as the template, with the m^3^C residue as the starting site of the extension reaction, which allows the direct view of the impacts of this modification to the whole reverse transcription complex. The reverse transcription yields or fidelity with different base pairing substrates in the presence of different reverse transcriptases were quantitated by the fluorescence gel images with single-nucleotide resolution (**Figure 4–6**) and the according UV images (**Figure S36-S38**).

**Figure 3.**
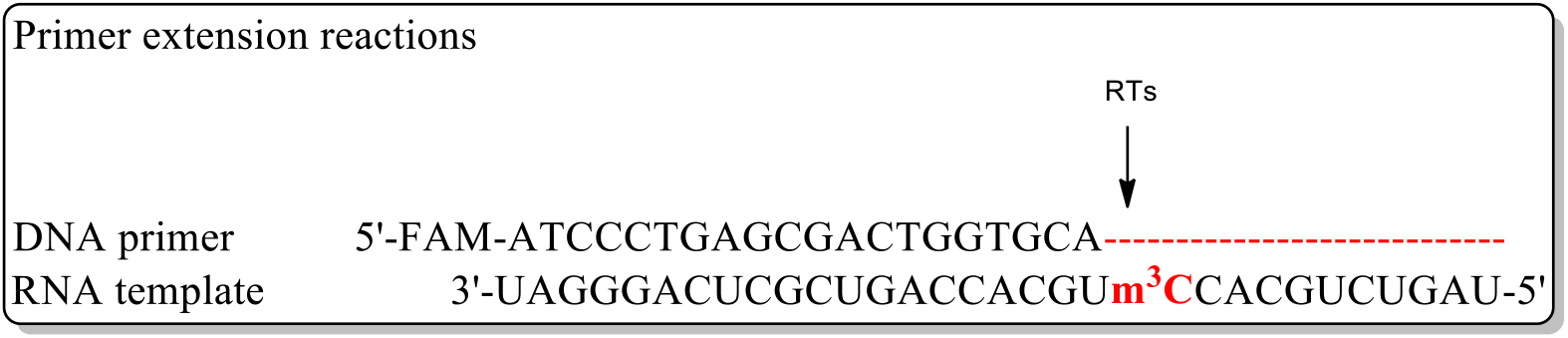
Primer extension reaction using m^3^C modified RNA template.

The Avian Myeloblastosis Virus Reverse Transcriptase (AMV-RT) is a widely used RNA-directed DNA polymerase in RT-PCR and RNA sequencing with high fidelity. (Myers et al., 1977) When AMV-RT was used in the presence of different dNTP substrates with native RNA template (**Figure 4A**), the reverse transcription reaction completes with all the natural dNTPs (lane Nat). AMV-RT could only use dGTP for incorporation against the starting C residue on the template, while no other dNTPs can be added to the primer (lane A, T, G, C). For AMV-RT and an m^3^C modified RNA template (**Figure 4B**), no full-length product was observed even in the presence of all the natural dNTPs (lane Nat *vs* N), indicating that this single m^3^C modification template completely inhibits the AMV-RT activity. Our observation is also consistent with the report that m^3^C acts as an RT stop residue in RT-based techniques.(Motorin et al., 2007)

**Figure 4.**
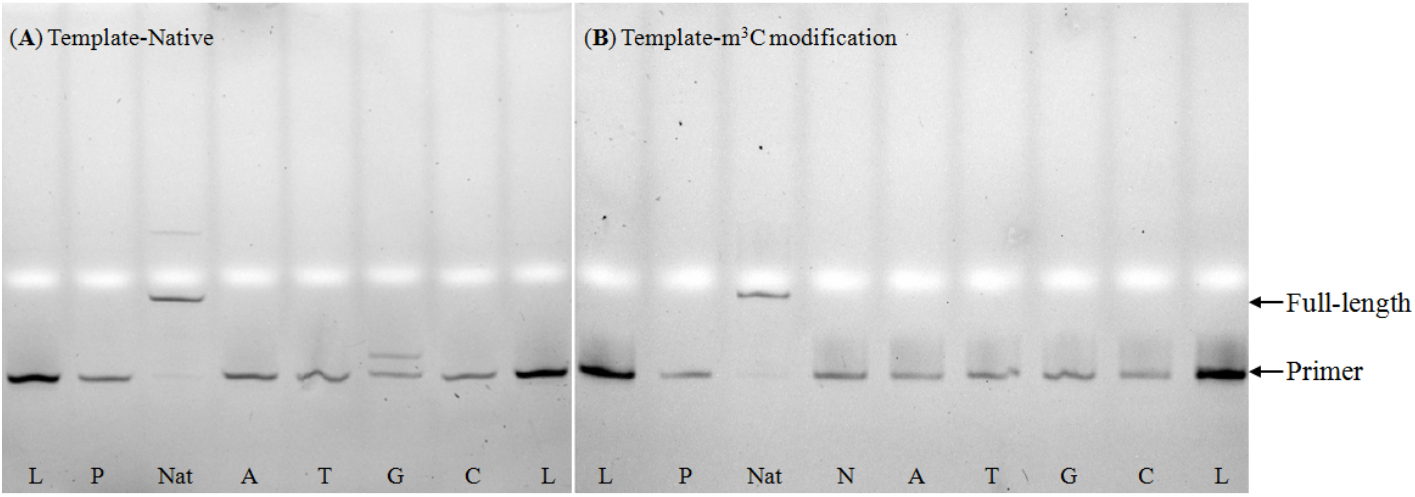
Fluorescent (**A and B**) gel images of standing-start primer extension reactions for AMV RT indicated using m^3^C containing RNA template and the corresponding natural template. Lanes: L, ladder; P, primer; Nat, natural template with all four dNTPs; A, T, G, and C, reactions in the presence of the respective dNTP; N, reactions in the presence of all four dNTPs.

By contrast, when the HIV-1-RT, which has relatively lower replication fidelity than AMV-RT, was used, it dramatically increased incorporation yield when dGTP was the lone dNTP (**Figure 5A**, lane G). We also observed that mis-incorporation of dATP and dTTP, but not dCTP, was present when using the HIV-1-RT (lane A, T and C). In the presence of m^3^C modified template (**Figure 5B**), we observed a very trace amount of full-length product (less than 5%) in the presence of all natural dNTPs (lanes Nat and N), supporting the idea that a single m^3^C modification severely inhibits the HIV-1-RT activity and results in a very low reaction yield. Interestingly, the m^3^C modification totally inhibits dGTP and dTTP incorporation but significantly increases the dATP incorporation yield (lanes A, T and G), resulting the order of preferential incorporation efficiency as A>>T>G, C.

**Figure 5.**
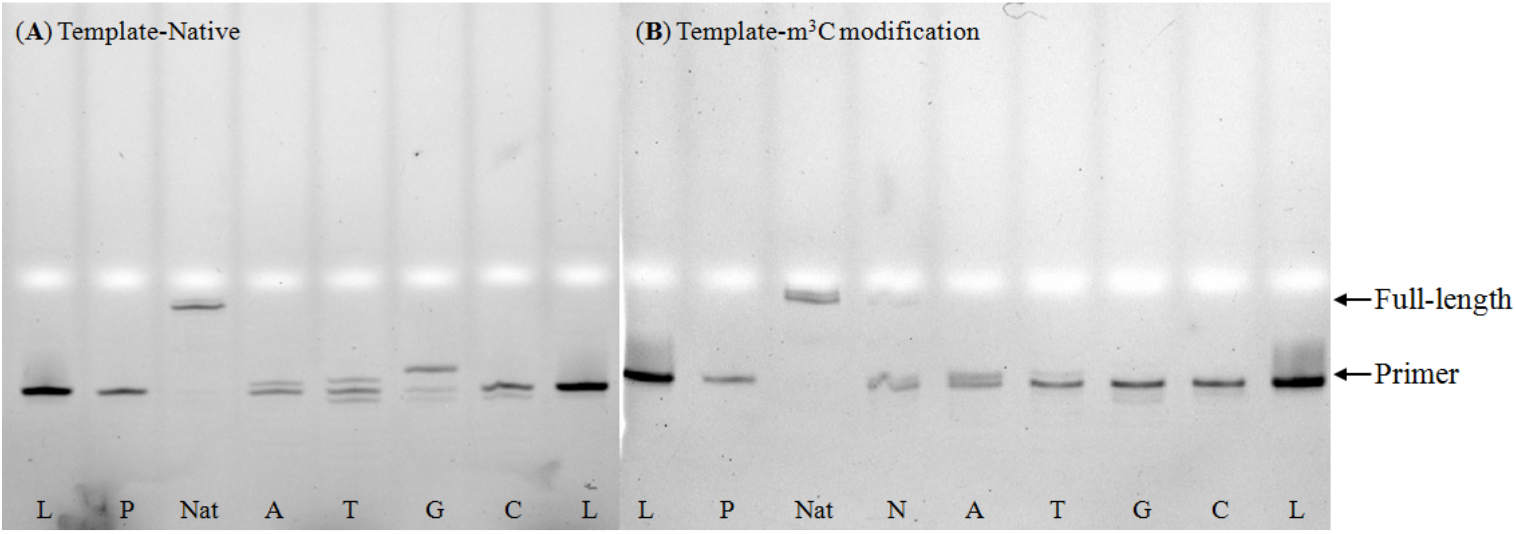
Fluorescent (**A and B**) gel images of standing-start primer extension reactions for HIV-1 RT indicated using m^3^C containing RNA template and the corresponding natural template. Lanes: L, ladder; P, primer; Nat, natural template with all four dNTPs; A, T, G, and C, reactions in the presence of the respective dNTP; N, reactions in the presence of all four dNTPs.

Subsequently, we explored the impacts of m^3^C on MMLV-RT, another enzyme with replication fidelity lower than AMV but higher than HIV-1 RT.(Skasko et al., 2005) MMLV-RT together with native RNA template (**Figure 6A**) in the reverse transcription reaction gives normal full length product in the presence of all the natural dNTPs (lane Nat). Although very high dGTP incorporation efficiency (lane G) was observed for MMLV-RT, the mis-incorporation of dATP was extremely high (lane A), supporting the idea that MMLV-RT can recognize and well accommodate the C:A pair in the reverse transcription complex. Similarly, dTTP was also incorporated into the primer strand with a medium yield (lane T). On the other hand, the presence of m^3^C in the template inhibits the normal enzyme activity of MMLV-RT (Lane N and G in **Figure 6B**), resulting in a mixture of three major strands in the reaction system, although the full-length product inhibition is much lower than the HIV-1 RT system. However, the m^3^C template modification does not impact the incorporation of dATP. As a result, a similar dNTPs incorporation efficiency order, A>>T, G>C, as HIV-1 RT, was observed for MMLV-RT. Furthermore, when the MutiScribe™ RT, a recombinant version of MMLV RT, was applied to the system, very similar results were obtained. (**Figure 6C** and **6D**).

**Figure 6.**
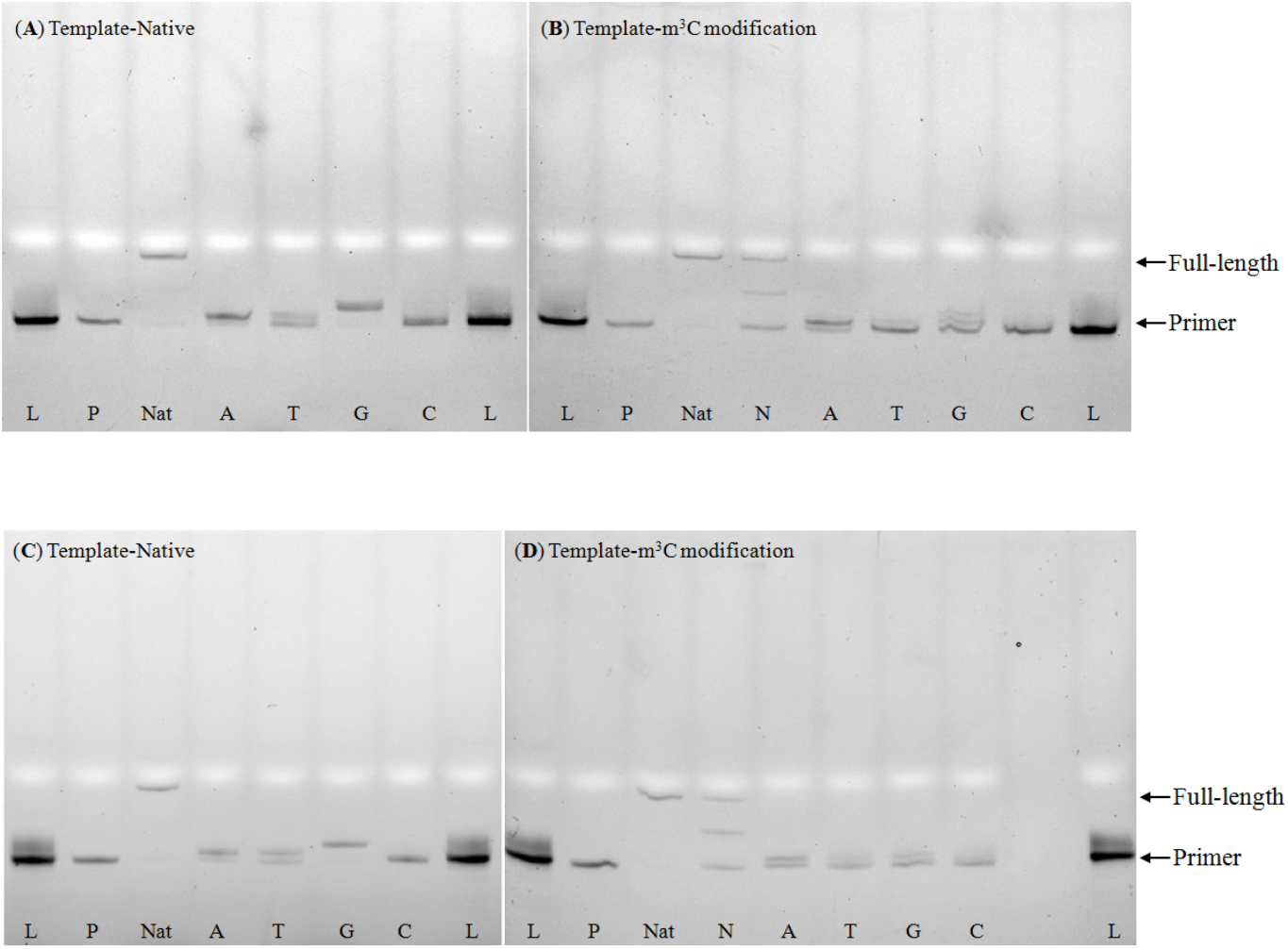
Fluorescent gel images of standing-start primer extension reactions for MMLV RT (**A** and **B**) and MultiScribe™ RT (**C** and **D**) indicated using m^3^C containing RNA templates and the corresponding natural templates. Lanes: L, ladders; P, primer; Nat, natural template with all four dNTPs; A, T, G, and C, reactions in the presence of the respective dNTP; N, reactions in the presence of all four dNTPs.

Many base modifications that do not change the Watson-Crick base pairing pattern have big impacts on the activity and fidelity of RNA polymerases, with m^6^A, m^5^C, m^5^U, hm^5^U being examples. (Potapov et al., 2018) The accuracy or fidelity of the base incorporation against a specific modified base on the template strand used for reverse transcription as well as the overall DNA or RNA synthesis error rates remains largely unestablished for many RNA modifications. HIV-1 RT is known as a low fidelity reverse transcriptase that catalyzes nucleotide mismatches with an error frequency of 1/2000 to 1/4000, and prefers a C:A pair over other mismatches, which frequently results in a G-to-A mutation during HIV gene replication.(Preston et al., 1988) Our results suggest that although the HIV-1 RT can induce both C:A and C:T pairs using an unmodified template, the presence of the m^3^C modification largely enhances the C:A pair while inhibiting the C:T one. We also observed that MMLV RT and MutiScribe™ RT cases enhances the C:A pair while inhibiting the C:T one. Our results support the idea that the m^3^C modification could specify the C:A pair in the presence of lower fidelity reverse transcriptases, thus further increasing the G-to-A mutation rate during reverse transcription. In contrast, the m^3^C encountered by enzymes with higher fidelity (i.e., AMV-RT) does not specify A or induce any other base mismatch, and primary serves as an RT stop.

We do note that the overall replication using HIV-1 RT is significantly (>90%) inhibited in the presence of m^3^C on the template, very similar to the AMV-RT case, which completely inhibits the DNA synthesis. One would expect a lower-fidelity RNA polymerase like HIV-1 to better accommodate the unnatural base pairs and promote higher replication yields when compared to the higher-fidelity AMV-RT do. Indeed, both MMLV RT and MutiScribe™ RT, which have a fidelity between HIV-1 RT and AMV-RT, show partial inhibition and result in a 50% of full-length product. Therefore, the m^3^C modification may have unique effects on HIV-1 RT, and allow the virus to regulate gene replication during different environmental selection stresses, which could be exploited to develop new therapeutics for HIV. Furthermore, the downstream demethylation process of m^3^C catalyzed by ALKBH3 may also play roles in restoring base pairing fidelity during virus replication, therefore representing another potential target in developing RNA based antiviral drugs.

## CONCLUSION

In summary, we synthesized the m^3^C phosphoramidite and a series of RNA oligonucleotides containing the modification. Our base-pairing and specificity studies show that the m^3^C modification disrupts the C:G pair and significantly decreases RNA duplex stability, which also results in the loss of base pairing discrimination of C:G pair with C:A, C:T, and C:C mismatched pairs. We also demonstrated that introducing two m^3^C modifications in the same sides (5’ or 3’ side) provided relative smaller effect on *T*_m_ compared to one modification. On the contrary, separating the two modifications could significantly reduce the duplex stability. Our molecular dynamic simulation study further reveals the detailed structural insights into the m^3^C:G base pairing pattern in RNA duplex. In addition, our investigation of this methylation effects on reverse transcription model demonstrated that the m^3^C modification could specify the C:A pair for some RT enzymes, which would induce the G to A mutation if used by low fidelity enzymes. For reverse transcriptase enzymes with higher fidelity (i.e., AMV-RT), m^3^C could completely shut down DNA synthesis. Our work provides detailed insights into the thermostability and the importance of m^3^C in RNA. Further it provides a foundation for exploiting the biochemical and biomedical potential of m^3^C in the design and development of RNA based therapeutics.

## MATERIALS AND METHODS

### Materials and general procedures of synthesis

Anhydrous solvents were used and redistilled using standard procedures. All solid reagents were dried under a high vacuum line prior to use. Air sensitive reactions were carried out under argon. RNase-free water, tips and tubes were used for RNA purification and thermodynamic studies. Analytical TLC plates pre-coated with silica gel F254 (Dynamic Adsorbents) were used for monitoring reactions and visualized by UV light. Flash column chromatography was performed using silica gel (32-63 μm). All ^1^H, ^13^C and ^31^P NMR spectra were recorded on a Bruker 400 and 500 MHz spectrometer. Chemical shift values are in ppm. ^13^C NMR signals were determined by using APT technique. High-resolution MS were achieved by ESI at University at Albany, SUNY.

### Synthesis of m^3^C phosphoramidites

#### 3-N-methyl-cytidine 2

To a solution of cytidine **1** (4.86 g, 20 mmol) in dry DMF (50 mL) was added iodomethane (2.5 mL, 40 mmol), and the solution was kept at room temperature during 24 hours. DMF was evaporated; the residue was evaporated with toluene (2 x 100 mL) and dissolved in acetone (20 mL). Hexane (50 mL) was added to the solution, and the resulting mixture was kept at −20 ^o^C for 1 hour. The precipitate was filtered, washed with cold mixture acetone:hexane (v/v = 1:1) (2 x 50 mL). The solid was dried in vacuum to give compound **2** (3.6 g, 14 mmol, 70% yield) as a yellow solid. ^1^H NMR (500 MHz, DVISO-*d*6) δ 9.79 (br, 1H), 9.15 (br, 1H), 8.31 (d, *J* = 7.5 Hz, 1H), 6.20 (d, *J* = 8.0 Hz, 1H), 5.70 (d, *J* = 3.5 Hz, 1H), 5.51 (br, 1H), 5.17 (br, 1H), 4.05-4.03 (m, 1H), 3.95-3.89 (m, 2H), 3.73 (dd, *J* = 2.5, 12.5 Hz, 1H), 3.60 (dd, *J* = 2.5, 12.0 Hz, 1H), 3.35 (s, 3H). ^13^C NMR (125 MHz, CDCl_3_) δ 159.4, 148.1, 142.1, 94.5, 91.2, 85.1, 74.6, 69.0, 60.2, 31.2. HRMS (ESI-TOF) [M+H]^+^ = 258.1090 (calc. 258.1090). Chemical formula: C_10_H_15_N_3_O_5_.

#### 1-(5’-O-4,4’-dimethoxytrityl-beta-D-ribofuranosyl)-3-N-methyl-cytidine 3

To a solution of compound **2** (3.5 g, 13.6 mmol) in dry pyridine (40 mL) was added 4,4’-dimethoxytrityl chloride (6.9 mg, 20.4 mmol). The resulting solution was stirred at RT overnight. To the suspension was added CH_2_Cl_2_ (200 mL) and the organic layer was subsequently washed with 5% aqueous Na_2_S_2_O_3_ (2 x 100 mL), sat. aqueous sodium bicarbonate (100 mL) and brine (100 mL). The organic layer was dried by anhydrous sodium sulfate, filtered and evaporated under reduced pressure. The residue was purified by silica gel chromatography to give compound **3** (3.9 g, 5.9 mmol, 43% yield) as a white solid. TLC R_*f*_ = 0.2 (10% MeOH in CH_2_Cl_2_). ^1^H NMR (400 MHz, CDCl_3_) δ 7.61(d, *J* = 8.0 Hz, 1H), 7.39-7.36 (m, 2H), 7.28-7.16 (m, 7H), 6.82-6.79 (m, 4H), 5.86 (d, *J* = 3.2 Hz, 1H), 5.76 (d, *J* = 7.2 Hz, 1H), 4.36-4.30 (m, 2H), 4.18 (m, 1H), 3.73 (d, 6H), 3.48-3.40 (m, 5H). ^13^C NMR (125 MHz, CDCl_3_) δ 158.9, 158.6, 149.4, 149.37, 144.5, 135.5, 135.3, 130.2, 128.2, 128.0, 113.3, 98.2, 90.9, 86.9, 83.5, 74.9, 69.9, 62.5, 55.3, 30.3. HRMS (ESI-TOF) [M+H]^+^ = 560.2390 (calc. 560.2397). Chemical formula: C_31_H_33_N_3_O_7_.

#### 1-(5’-O-4,4’-dimethoxytrityl-beta-D-ribofuranosyl)-4-N-benzoyl-3-N-methyl-cytidine 4

Compound **3** (2.3 g, 4.2 mmol) was co-evaporated with pyridine (2 x 50 mL) and redissolved in pyridine (50 ml). Trimethylsilyl chloride (TMSCl) (2.1 mL, 16.8 mmol) was added and the mixture was stirred at RT for 1 h whereupon benzoyl chloride (BzCl) (0.84 mL, 5.04 mmol) was added. The resulting solution was stirred for 4 h at RT whereupon water (10 mL) was added. After stirring for 5 min at RT, aqueous ammonia (15 mL, 15.8 M) was added and the mixture was stirred for 15 min at RT and then evaporated to dryness under reduced pressure. The residue was purified by silica gel chromatography to give compound **4** (2.3 g, 3.47 mmol, 82% yield) as a white solid. TLC R_*f*_ = 0.4 (50% EtOAc in CH_2_Cl_2_). ^1^H NMR (400 MHz, CDCl_3_) δ 8.15-8.12 (m, 2H), 7.64 (d, *J*= 8.4 Hz, 1H),7.56-7.51 (m, 1H), 7.46-7.42 (m, 2H), 7.35-7.33 (m, 2H), 7.29-7.18 (m, 8H), 6.84-6.80 (m, 4H), 6.25 (d, *J* = 8.4 Hz, 1H), 5.82 (d, *J* = 3.6 Hz, 1H), 4.37-4.26 (m, 3H), 3.78 (d, 6H), 3.58 (s, 3H). 3.48 (dd, *J* = 2.8 Hz, 10.8 Hz, 1H), 3.38 (dd, *J* = 3.2 Hz, 10.8 Hz, 1H). ^13^C NMR (125 MHz, CDCl_3_) δ 177.4, 158.7, 151.1, 144.1, 135.8, 135.7, 135.4, 135.2, 132.5, 130.1, 130.0, 129.7, 128.2, 128.1, 128.0, 127.1, 113.31, 113.30, 98.2, 91.5, 87.1, 84.3, 76.2, 70.5, 62.2, 55.2, 30.0. HRMS (ESI-TOF) [M+H]^+^ = 664.2646 (calc. 664.2659). Chemical formula: C_38_H_37_N_3_O_8_.

#### 1-(2’-O-tert-butyldimethylsilyl-5’-O-4,4’-dimethoxytrityl-beta-D-ribofuranosyl)-4-N-benzoyl-3-N-methyl-cytidine 5

Compound **4** (1.3 g, 2 mmol) was dissolved in dry DMF (12 mL), then tert-butyldimethylsilyl chloride (TBDMSCl, 362 mg, 2.4 mmol) and imidazole (272 mg, 4 mmol) were added into the solution. The resulting solution was stirred overnight at RT. The solution was diluted with EtOAc (200 mL) and washed with brine (2 x 100 mL). The organic layer was dried by anhydrous sodium sulfate, filtered and evaporated under reduced pressure. The residue was purified by silica gel chromatography to give compound **5** (600 mg, 0.77 mmol, 39% yield) as a white solid. TLC R_*f*_ = 0.6 (Hexane:EA = 1:1). ^1^H NMR (500 MHz, CDCl_3_) δ 8.15-8.12 (m, 2H), 7.82 (d, *J* = 8.5 Hz, 1H), 7.55-7.50 (m, 1H), 7.46-7.42 (m, 2H), 7.38-7.35 (m, 2H), 7.29-7.25 (m, 6H), 7.22-7.16 (m, 1H), 6.85-6.82 (m, 4H), 6.08 (d, *J* = 8.0 Hz, 1H), 5.94 (d, *J* = 2.5 Hz, 1H), 4.38-4.33 (m, 1H), 4.29-4.27 (m, 1H), 4.09-4.06 (m, 1H), 3.78 (d, *J* = 1.0 Hz, 6H), 3.55 (s, 3H), 3.57-3.46 (m, 2H), 0.94 (s, 9H), 0.24 (s, 3H), 0.18 (s, 3H). ^13^C NMR (125 MHz, CDCl_3_) δ 177.3, 158.7, 155.7, 150.3, 144.1, 136.0, 135.9, 135.4, 135.2, 132.4, 130.1, 130.0, 129.7, 128.20, 128.19, 128.0, 127.2, 98.1, 89.8, 87.1, 83.3, 76.8, 69.8, 61.9, 55.2, 30.0, 25.8, 18.1, −4.5, −5.2. HRMS (ESI-TOF) [M+H]^+^ = 778.3527 (calc. 778.3524). Chemical formula: C_44_H_51_N_3_O_8_Si.

#### 1-[2’-O-tert-butyldimethylsilyl-3’-O-(2-cyanoethyl-N,N-diisopropylamino)phosphor amidite-5’-O-(4,4’-dimethoxytrityl-beta-D-ribofuranosyl)]-4-N-benzoyl-3-N-methyl-cytidine 6

To a solution of compound **5** (600 mg, 0.77 mmol) in dry DCM (13 mL) was added N,N-di-iso-propylethylamine (0.38 mL, 3.08 mmol) and 2-cyanoethyl *N,N*-diisopropylchlorophosphoramidite (0.23 mL, 1.54 mmol). The resulting solution was stirred overnight at room temperature under argon gas. The reaction was quenched with water and extracted with ethyl acetate. After drying the organic layer over Na_2_SO_4_ and evaporation. The residue was purified by silica gel chromatography to give compound **6** (500 mg, 0.51 mmol, 66% yield) as a white solid. TLC R_*f*_ = 0.6 (Hexane:EA = 1:1). ^1^H NMR (400 MHz, CDCl_3_) δ 8.15-8.12 (m, 2H), 7.90-7.19 (m, 13H), 6.84-6.80 (m, 4H), 6.16-5.91 (m, 2H), 4.38-3.91 (m, 3H), 3.78 (s, 6H), 3.61-3.36 (m, 7H), 2.67-2.37 (m, 2H), 2.06-2.04 (m, 1H), 1.29-1.14 (m, 12H), 1.02-0.98 (m, 3H), 0.93-0.9 (m, 9H), 0.19-0.13 (m, 6H). ^31^P NMR (162 MHz, CDCl_3_) δ 149.97, 148.94. HRMS (ESI-TOF) [M+H]^+^ = 978.4561 (calc. 978.4602). Chemical formula: C_53_H_68_N_5_O_9_PSi.

### Synthesis and purification of m^3^C containing RNA oligonucleotides

All oligonucleotides were chemically synthesized at 1.0 μmol scales by solid phase synthesis using the Oligo-800 synthesizer. The m^3^C phosphoramidite was dissolved in acetonitrile to a concentration of 0.1 M. I2 (0.02 M) in THF/Py/H_2_O solution was used as an oxidizing reagent. Coupling was carried out using 5-ethylthio-1H-tetrazole solution (0.25 M) in acetonitrile for 12 min, for both native and modified phosphoramidites. About 3% trichloroacetic acid in methylene chloride was used for the 5’-detritylation. Synthesis was performed on control-pore glass (CPG-500) immobilized with the appropriate nucleoside through a succinate linker. All the reagents used are standard solutions obtained from ChemGenes Corporation. The oligonucleotide was prepared in DMTr off form. After synthesis, the oligos were cleaved from the solid support and fully deprotected with 1:1 v/v ammonium hydroxide solution (28% NH_3_ in H_2_O) and Methylamine (40% w/w aqueous solution) at 65 °C for 45 min. The solution was evaporated to dryness by Speed-Vac concentrator. The solid was dissolved in 100 μL DMSO and was desilylated using a triethylamine trihydrogen fluoride (Et_3_N•3HF) solution at 65 °C for 2.5 h. Cooled down to room temperature the RNA was precipitated by adding 0.025 mL of 3 M sodium acetate and 1 mL of ethanol. The solution was cooled to −80 °C for 1 h before the RNA was recovered by centrifugation and finally dried under vacuum.

The oligonucleotides were purified by IE-HPLC at a flow rate of 1 mL/min. Buffer A was 20 mM Tris-HCl, pH 8.0; buffer B 1.25M NaCl in 20 mM Tris-HCl, pH 8.0. A linear gradient from 100% buffer A to 70% buffer B in 20 min was used to elute the oligos. The analysis was carried out by using the same type of analytical column with the same eluent gradient. All the modified-oligos were checked by MALDI MS. The 22-mer and 31-mer RNA oligonucleotides were purified on a preparative 20% denaturing polyacrylamide gel (PAGE).

### UV-melting temperature (*T*_m_) study

Solutions of the duplex RNAs (1.5 μM) were prepared by dissolving the purified RNAs in sodium phosphate (10 mM, pH 7.0) buffer containing 100 mM NaCl. The solutions were heated to 95 °C for 5 min, then cooled down slowly to room temperature, and stored at 4 °C for 2 h before *T*_m_ measurement. Thermal denaturation was performed in a Cary 300 UV-Visible Spectrophotometer with a temperature controller. The temperature reported is the block temperature. Each denaturizing curve was acquired at 260 nm by heating and cooling from 5 to 80 °C for four times at a rate of 0.5 °C/min. All the melting curves were repeated at least four times. The thermodynamic parameter of each strand was obtained by fitting the melting curves using the Meltwin software.

### Molecular dynamic simulation studies

To study the m^3^C modification in the context of the RNA duplex in MD simulations, we developed AMBER (Cornell et al., 1995) type force-field parameters for the atoms of the modified nucleoside. We used the AM1-BCC (Jakalian et al., 2002) charge model to calculate the atomic charges, which is developed as a fast yet accurate alternate for ESP-fit using Hartree-Fock theory and 6-31G* basis-sets (Cornell et al., 1993). AMBER99 force-field parameters were used for bonded interactions (Cornell et al., 1995), and AMBER99 parameters with Chen-Garcia corrections (Chen and Garcia, 2013) for the bases were used for LJ interactions. The unmodified RNA duplex was constructed in a-form using the Nucleic Acid Builder (NAB) suite of AMBER, and mutated to create the modification.

Molecular dynamics simulations were performed using Gromacs-2018 package (Abraham et al., 2015). The simulation system included the RNA duplex in water in a 3D periodic box. The initial box size was 4.0 x 4.0 x 6.0 nm^3^ containing the RNA duplex, 3060 water molecules, and 22 neutralizing Na^+^ ions. The system was subjected to energy minimization to prevent any overlap of atoms, followed by a 1 ns equilibration run. The equilibrated system was then subjected to a 100 ns production run. The MD simulations incorporated leap-frog algorithm with a 2 fs timestep to integrate the equations of motion. The system was maintained at 300K and 1 bar, using the velocity rescaling thermostat (Bussi et al., 2007) and Parrinello-Rahman barostat (Berendsen et al., 1984), respectively. The long-ranged electrostatic interactions were calculated using particle mesh Ewald (PME) (Darden et al., 1993) algorithm with a real space cut-off of 1.2 nm. LJ interactions were also truncated at 1.2 nm. TIP3P model (Jorgensen et al., 1983) was used represent the water molecules, and LINCS (Hess et al., 1997) algorithm was used to constrain the motion of hydrogen atoms bonded to heavy atoms. Co-ordinates of the RNA molecule were stored every 20 ps for further analysis.

### Reverse transcription (RT) assays

RT assays were performed with AMV RT (ThermoFisher), HIV-1 RT (AS ONE Corp.), MMLV RT (ThermoFisher) and MutiScribe™ RT (ThermoFisher) in 20 μL total solution containing 10X reverse transcription buffer: 50 mM Tris (pH 8.3), 75 mM KCl, 3 mM MgCl_2_, 10 mM DTT. Final reaction mixtures contained RNA template (5 μM), DNA FAM-primer (2.5 μM) and dNTP (1 mM). After addition of Rnase inhibitor (20 U) and each RTs: AMV RT (10 U), HIV-1 RT (4 U), MMLV (100 U) and MutiScribe™ (50 U), the mixtures were incubated at 37 °C for 1 h. The reactions were quenched with stop solution [98% formamide, 0.05% xylene cyanol (FF), and 0.05% bromophenol blue], heated to 90 °C for 5 min and then cooled to 0 °C at ice-bath. Reactions were analyzed by 15% PAGE 8 M urea at 250 V for 1-1.5 h. The fluorescent and UV gel imaging were done on a Bio-Rad Gel XR+ imager.

## ASSOCIATED CONTENT

### Supporting Information

Electronic Supporting Information (ESI) available: Experimental procedures, spectral data, UV-melting curves and PAGE gel UV-images.

## ACKNOWLEDGMENTS

We are grateful to NSF (CHE-1845486) and NERF grant from the University at Albany, State University of New York for the financial support. We thank Drs. Zhen Huang and Cen Chen for their help in MS-Spec experiments.

## REFERENCES

Abraham, M.J., Murtola, T., Schulz, R., Páll, S., Smith, J.C., Hess, B., and Lindahl, E. (2015). GROMACS: High performance molecular simulations through multi-level parallelism from laptops to supercomputers. SoftwareX 1-2, 19–25.

Basanta-Sanchez, M., Temple, S., Ansari, S.A., D’Amico, A., and Agris, P.F. (2016). Attomole quantification and global profile of RNA modifications: Epitranscriptome of human neural stem cells. Nucleic Acids Res. 44, e26.

Berendsen, H.J.C., Postma, J.P.M., van Gunsteren, W.F., DiNola, A., and Haak, J.R. (1984). Molecular dynamics with coupling to an external bath. J. Chem. Phys. 81, 3684–3690.

Boccaletto, P., Machnicka, M.A., Purta, E., Piatkowski, P., Baginski, B., Wirecki, T.K., de Crecy-Lagard, V., Ross, R., Limbach, P.A., Kotter, A., et al. (2018). MODOMICS: a database of RNA modification pathways. 2017 update. Nucleic Acids Res. 46, D303–D307.

Brookes, P., and Lawley, P.D. (1962). The Methylation of Cytosine and Cytidine. J. Chem. Soc. 1348–1351.

Bussi, G., Donadio, D., and Parrinello, M. (2007). Canonical sampling through velocity rescaling. J. Chem. Phys. 126, 014101.

Cantara, W.A., Crain, P.F., Rozenski, J., McCloskey, J.A., Harris, K.A., Zhang, X., Vendeix, F.A., Fabris, D., and Agris, P.F. (2011). The RNA Modification Database, RNAMDB: 2011 update. Nucleic Acids Res. 39, D195–201.

Chen, A.A., and Garcia, A.E. (2013). High-resolution reversible folding of hyperstable RNA tetraloops using molecular dynamics simulations. Proc. Natl. Acad. Sci. U S A 110, 16820–16825.

Chen, B., Li, Y., Song, R.F., Xue, C., and Xu, F. (2019a). Functions of RNA N6-methyladenosine modification in cancer progression. Mol. Biol. Rep. 46, 1383–1391.

Chen, Z., Qi, M., Shen, B., Luo, G., Wu, Y., Li, J., Lu, Z., Zheng, Z., Dai, Q., and Wang, H. (2019b). Transfer RNA demethylase ALKBH3 promotes cancer progression via induction of tRNA-derived small RNAs. Nucleic Acids Res. 47, 2533–2545.

Ciuffi, A. (2016). Viral cell biology: HIV RNA gets methylated. Nat. Microbiol. 1, 16037.

Clark, W.C., Evans, M.E., Dominissini, D., Zheng, G., and Pan, T. (2016). tRNA base methylation identification and quantification via high-throughput sequencing. Rna 22, 1771–1784.

Cornell, W.D., Cieplak, P., Bayly, C.I., Gould, I.R., Merz, K.M., Ferguson, D.M., Spellmeyer, D.C., Fox, T., Caldwell, J.W., and Kollman, P.A. (1995). A Second Generation Force Field for the Simulation of Proteins, Nucleic Acids, and Organic Molecules. J. Am. Chem. Soc. 117, 5179–5197.

Cornell, W.D., Cieplak, P., Bayly, C.I., and Kollman, P.A. (1993). Application of RESP charges to calculate conformational energies, hydrogen bond energies, and free energies of solvation. J. Am. Chem. Soc. 115, 9620–9631.

Cozen, A.E., Quartley, E., Holmes, A.D., Hrabeta-Robinson, E., Phizicky, E.M., and Lowe, T.M. (2015). ARM-seq: AlkB-facilitated RNA methylation sequencing reveals a complex landscape of modified tRNA fragments. Nat. Methods 12, 879–884.

D’Silva, S., Haider, S.J., and Phizicky, E.M. (2011). A domain of the actin binding protein Abp140 is the yeast methyltransferase responsible for 3-methylcytidine modification in the tRNA anti-codon loop. Rna 17, 1100–1110.

Darden, T., York, D., and Pedersen, L. (1993). Particle mesh Ewald: An Nolog(N) method for Ewald sums in large systems. J. Chem. Phys. 98, 10089–10092.

Desrosiers, R., Friderici, K., and Rottman, F (1974). Identification of methylated nucleosides in messenger RNA from Novikoff hepatoma cells. Proc. Natl. Acad. Sci. U S A 71, 3971–3975.

Fu, Y., Dominissini, D., Rechavi, G., and He, C. (2014). Gene expression regulation mediated through reversible m(6)A RNA methylation. Nat. Rev. Genet. 15, 293–306.

Hall, R.H. (1963). Isolation of 3-Methyluridine and 3-Methylcytidine from Solubleribonucleic Acid. Biochem. Biophys. Res. Commun. 12, 361–364.

Han, L., Marcus, E., D’Silva, S., and Phizicky, E.M. (2017). S. cerevisiae Trm140 has two recognition modes for 3-methylcytidine modification of the anticodon loop of tRNA substrates. Rna 23, 406–419.

Hess, B., Bekker, H., Berendsen, H.J.C., and Fraaije, J.G.E.M. (1997). LINCS: A linear constraint solver for molecular simulations. J. Comput. Chem. 18, 1463–1472.

Holley, R.W., Apgar, J., Everett, G.A., Madison, J.T., Marquisee, M., Merrill, S.H., Penswick, J.R., and Zamir, A. (1965a). Structure of a Ribonucleic Acid. Science 147, 1462–1465.

Holley, R.W., Everett, G.A., Madison, J.T., and Zamir, A. (1965b). Nucleotide Sequences in the Yeast Alanine Transfer Ribonucleic Acid. J. Biol. Chem. 240, 2122–2128.

Hori, H. (2014). Methylated nucleosides in tRNA and tRNA methyltransferases. Front. Genet. 5, 144.

Iwanami, Y., and Brown, G.M. (1968). Methylated bases of transfer ribonucleic acid from HeLa and L cells. Arch. Biochem. Biophys. 124, 472–482.

Jakalian, A., Jack, D.B., and Bayly, C.I. (2002). Fast, efficient generation of high-quality atomic charges. AM1-BCC model: II. Parameterization and validation. J. Comput. Chem. 23, 1623–1641.

Jiang, Q., Crews, L.A., Holm, F., and Jamieson, C.H.M. (2017). RNA editing-dependent epitranscriptome diversity in cancer stem cells. Nat. Rev. Cancer 17, 381–392.

Jorgensen, W.L., Chandrasekhar, J., and Madura, J.D. (1983). Comparison of simple potential functions for simulating liquid water. J. Chem. Phys. 79, 926–935.

Lichinchi, G., Zhao, B.S., Wu, Y., Lu, Z., Qin, Y., He, C., and Rana, T.M. (2016). Dynamics of Human and Viral RNA Methylation during Zika Virus Infection. Cell Host Microbe 20, 666–673.

Liu, F., and He, C. (2017). A new modification for mammalian messenger RNA. J. Biol. Chem. 292, 14704–14705.

Machnicka, M.A., Milanowska, K., Osman Oglou, O., Purta, E., Kurkowska, M., Olchowik, A., Januszewski, W., Kalinowski, S., Dunin-Horkawicz, S., Rother, K.M., et al. (2013). MODOMICS: a database of RNA modification pathways--2013 update. Nucleic Acids Res. 41, D262–267.

McDowell, J.A., and Turner, D.H. (1996). Investigation of the structural basis for thermodynamic stabilities of tandem GU mismatches: solution structure of (rGAGGUCUC)2 by two-dimensional NMR and simulated annealing. Biochemistry 35, 14077–14089.

McIntyre, W., Netzband, R., Bonenfant, G., Biegel, J.M., Miller, C., Fuchs, G., Henderson, E., Arra, M., Canki, M., Fabris, D., et al. (2018). Positive-sense RNA viruses reveal the complexity and dynamics of the cellular and viral epitranscriptomes during infection. Nucleic Acids Res. 46, 5776–5791.

Mongan, N.P., Emes, R.D., and Archer, N. (2019). Detection and analysis of RNA methylation. F1000Research 8.

Motorin, Y., Muller, S., Behm-Ansmant, I., and Branlant, C. (2007). Identification of modified residues in RNAs by reverse transcription-based methods. Meth. Enzymol. 425, 21–53.

Myers, J.C., Spiegelman, S., and Kacian, D.L. (1977). Synthesis of full-length DNA copies of avian myeloblastosis virus RNA in high yields. Proc. Natl. Acad. Sci. U S A 74, 2840–2843.

Nachtergaele, S., and He, C. (2017). The emerging biology of RNA post-transcriptional modifications. RNA Biol. 14, 156–163.

Nachtergaele, S., and He, C. (2018). Chemical Modifications in the Life of an mRNA Transcript. Annu. Rev. Genet. 52, 349–372.

Noma, A., Yi, S., Katoh, T., Takai, Y., Suzuki, T., and Suzuki, T. (2011). Actin-binding protein ABP140 is a methyltransferase for 3-methylcytidine at position 32 of tRNAs in Saccharomyces cerevisiae. Rna 17, 1111–1119.

Ogilvie, K.K., and Kader, H.A. (1983). Synthesis of 3, N4–Dimethylcytidine. Nucleosides Nucleotides Nucleic Acids 2, 345–350.

Olson, M.V., Page, G.S., Sentenac, A., Piper, P.W., Worthington, M., Weiss, R.B., and Hall, B.D. (1981). Only one of two closely related yeast suppressor tRNA genes contains an intervening sequence. Nature 291, 464–469.

Ougland, R., Zhang, C.M., Liiv, A., Johansen, R.F., Seeberg, E., Hou, Y.M., Remme, J., and Falnes, P.O. (2004). AlkB restores the biological function of mRNA and tRNA inactivated by chemical methylation. Mol. Cell 16, 107–116.

Potapov, V., Fu, X., Dai, N., Correa, I.R., Jr., Tanner, N.A., and Ong, J.L. (2018). Base modifications affecting RNA polymerase and reverse transcriptase fidelity. Nucleic Acids Res. 46, 5753–5763.

Preston, B.D., Poiesz, B.J., and Loeb, L.A. (1988). Fidelity of HIV-1 reverse transcriptase. Science 242, 1168–1171.

Roundtree, I.A., Evans, M.E., Pan, T., and He, C. (2017). Dynamic RNA Modifications in Gene Expression Regulation. Cell 169, 1187–1200.

Sergiev, P.V., Aleksashin, N.A., Chugunova, A.A., Polikanov, Y.S., and Dontsova, O.A. (2018). Structural and evolutionary insights into ribosomal RNA methylation. Nat. Chem. Biol. 14, 226–235.

Shi, H., Wei, J., and He, C. (2019). Where, When, and How: Context-Dependent Functions of RNA Methylation Writers, Readers, and Erasers. Mol. Cell 74, 640–650.

Skasko, M., Weiss, K.K., Reynolds, H.M., Jamburuthugoda, V., Lee, K., and Kim, B. (2005). Mechanistic differences in RNA-dependent DNA polymerization and fidelity between murine leukemia virus and HIV-1 reverse transcriptases. J. Biol. Chem. 280, 12190–12200.

Song, J., and Yi, C. (2017). Chemical Modifications to RNA: A New Layer of Gene Expression Regulation. ACS Chem. Biol. 12, 316–325.

Ueda, Y., Ooshio, I., Fusamae, Y., Kitae, K., Kawaguchi, M., Jingushi, K., Hase, H., Harada, K., Hirata, K., and Tsujikawa, K. (2017). AlkB homolog 3-mediated tRNA demethylation promotes protein synthesis in cancer cells. Sci. Rep. 7, 42271.

Wang, X., Lu, Z., Gomez, A., Hon, G.C., Yue, Y., Han, D., Fu, Y., Parisien, M., Dai, Q., Jia, G., et al. (2014). N6-methyladenosine-dependent regulation of messenger RNA stability. Nature 505, 117–120.

Wu, L. (2019). HIV Evades Immune Surveillance by Methylation of Viral RNA. Biochemistry 58, 1699–1700.

Wu, Y., Tang, Y., Dong, X., Zheng, Y.Y., Haruehanroengra, P., Mao, S., Lin, Q., and Sheng, J. (2020). RNA Phosphorothioate Modification in Prokaryotes and Eukaryotes. ACS Chem. Biol. 15, 1301–1305.

Xu, L., Liu, X., Sheng, N., Oo, K.S., Liang, J., Chionh, Y.H., Xu, J., Ye, F., Gao, Y.G., Dedon, P.C., et al. (2017). Three distinct 3-methylcytidine (m(3)C) methyltransferases modify tRNA and mRNA in mice and humans. J. Biol. Chem. 292, 14695–14703.

Zaccara, S., Ries, R.J., and Jaffrey, S.R. (2019). Reading, writing and erasing mRNA methylation. Nat. Rev. Mol. Cell Bio. 20, 608–624.

