## supplemental Table 3, supplemental Figure 4-6 for "Base Pairing and Functional Insights into *N^3^*-methylcytidine (m^3^C) in RNA"

Table of Contents

**Part I** $^1$H and $^{13}$C NMR and HRMS spectra of synthesized compounds................................. S2-S9

**Part II** Synthesis, HPLC and Characterization of modified oligonucleotides.......................S9-S14

**Part III** UV-melting temperature ($T_m$) study……………………………………………...…....S14

**Part IV**. UV gel images of standing-start primer extension reactions S14-15

**Part I. $^{1}$H and $^{13}$C NMR and HRMS spectra of synthesized compounds**

**$^{1}$HNMR, $^{13}$CNMR and Mass Spectrum**

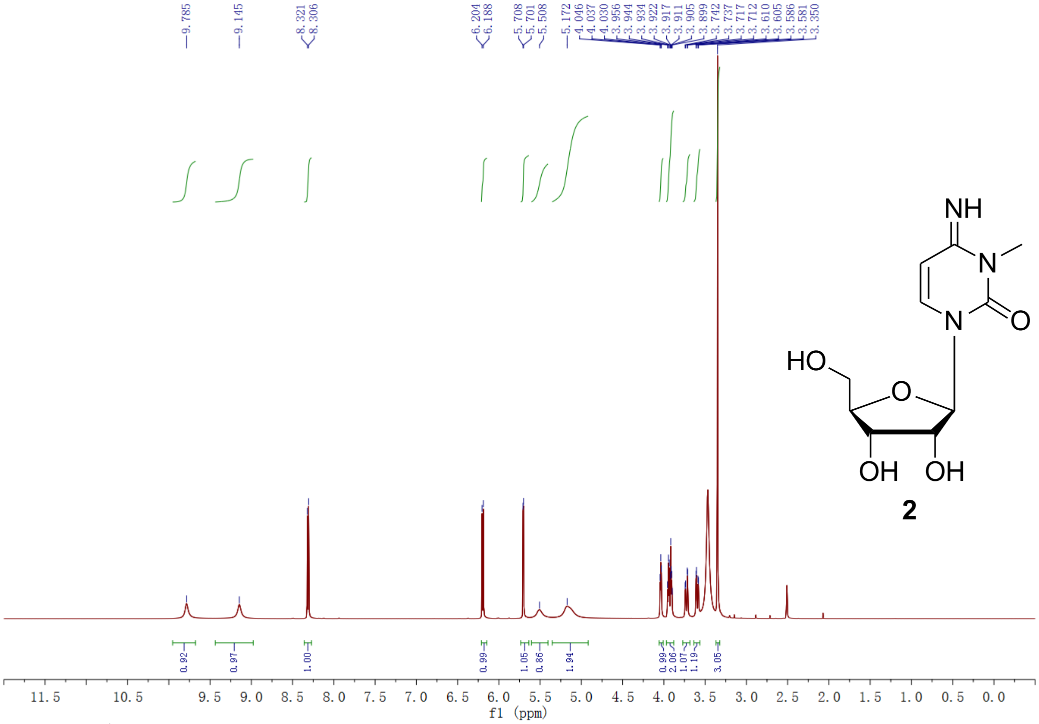


**Fig. S1.** $^{1}$H NMR of compound **2**.

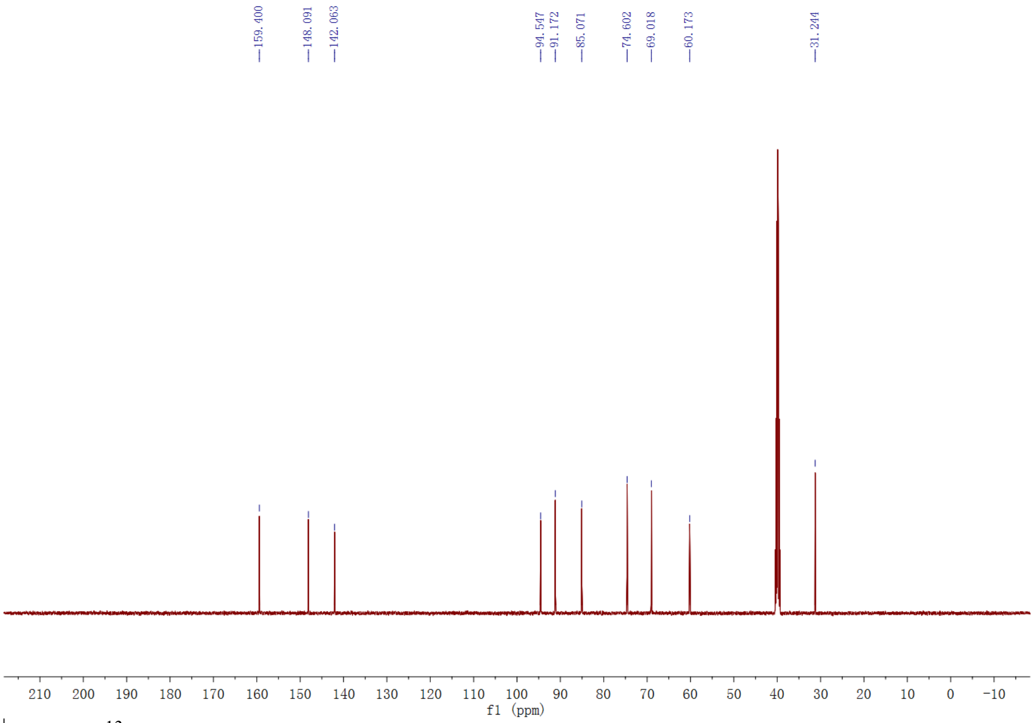


**Fig. S2.** $^{13}$C NMR of compound **2**.

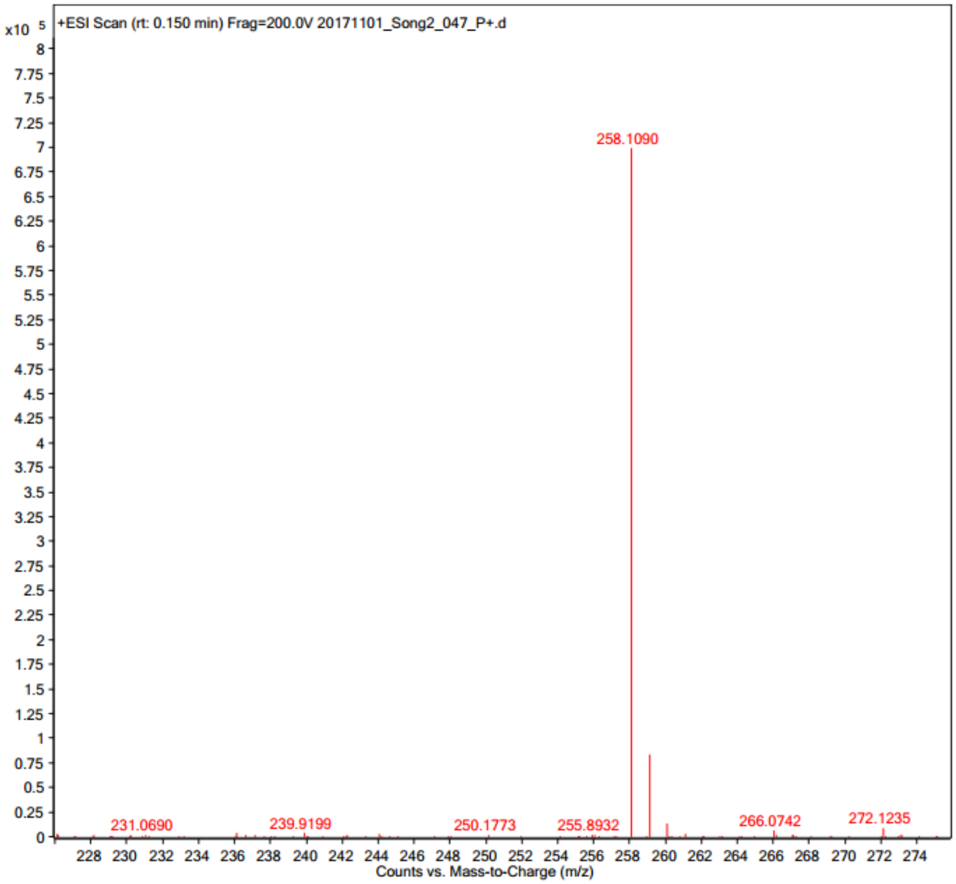


**Fig. S3.** Mass of compound **2**.

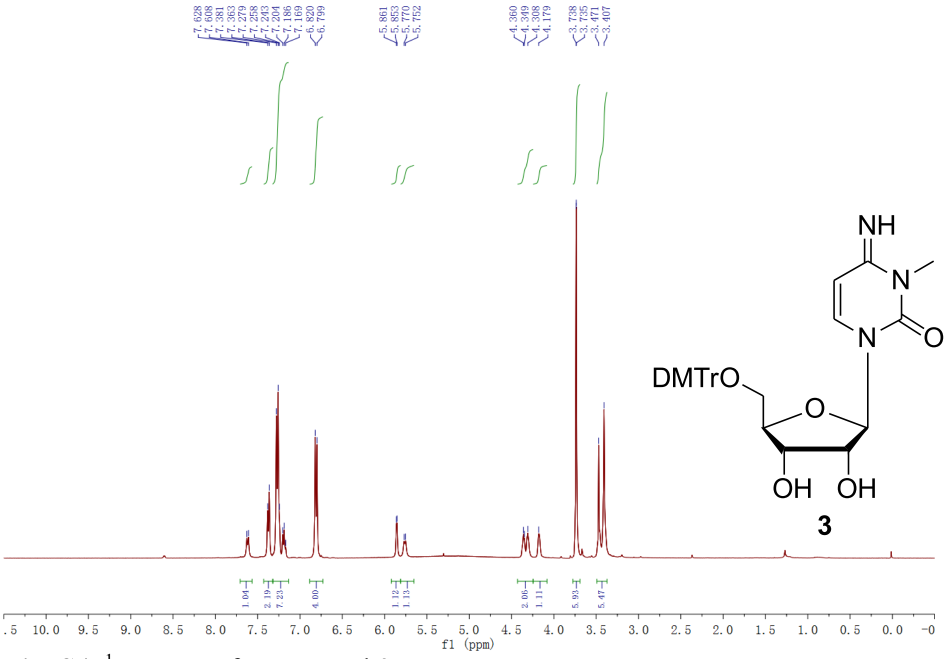


**Fig. S4.** $^{1}$H NMR of compound **3**.

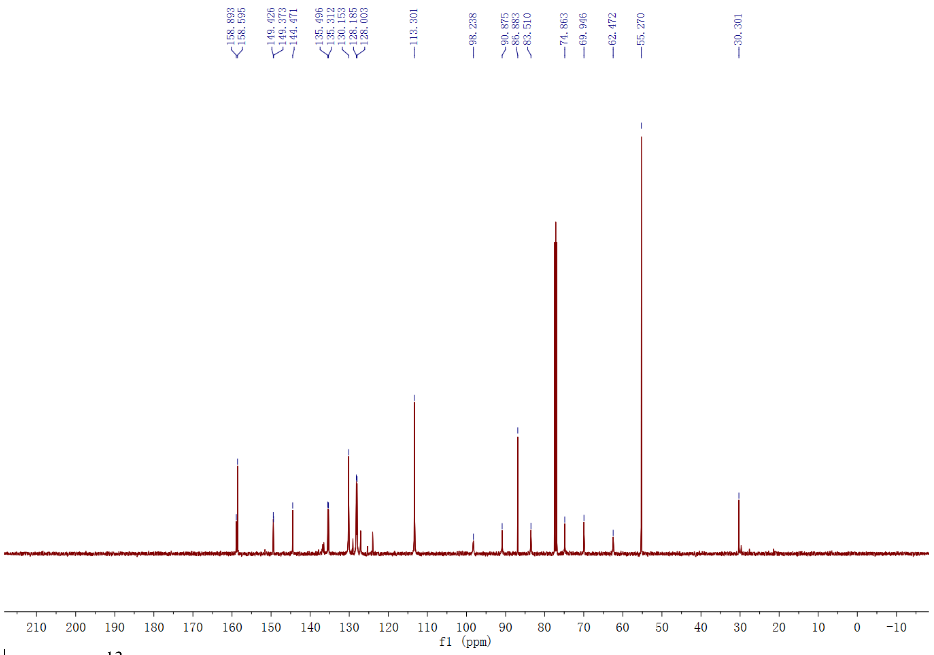


**Fig. S5.** $^{13}C$ NMR of compound **3**.

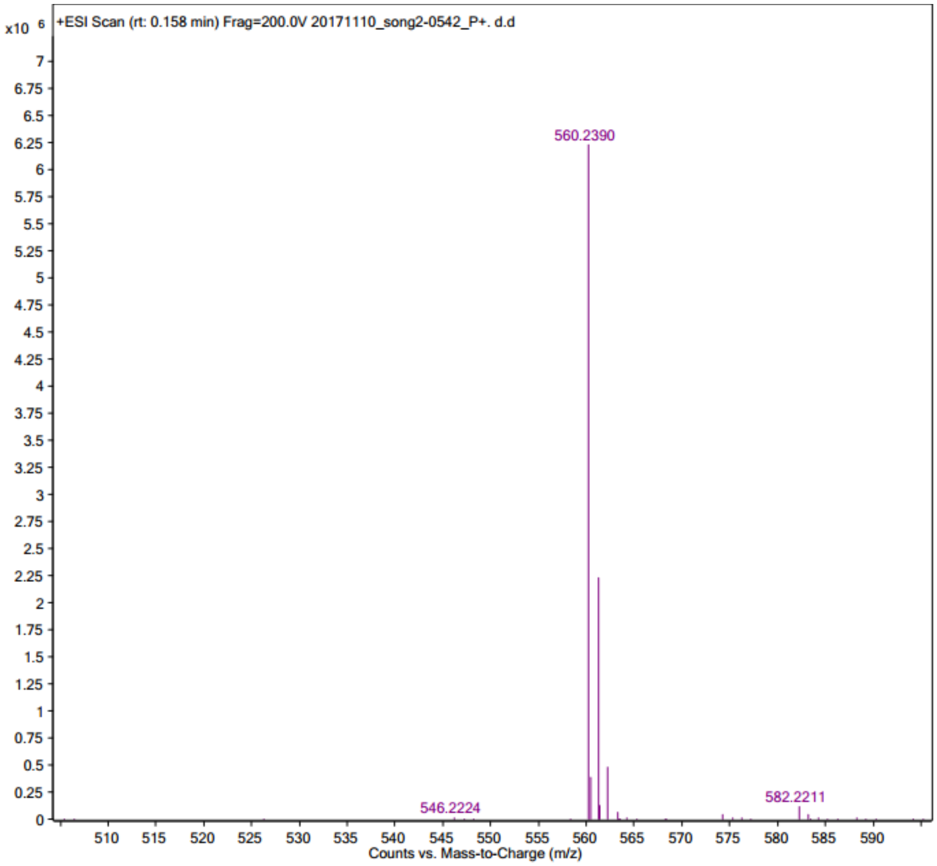


**Fig. S6.** Mass of compound **3**.

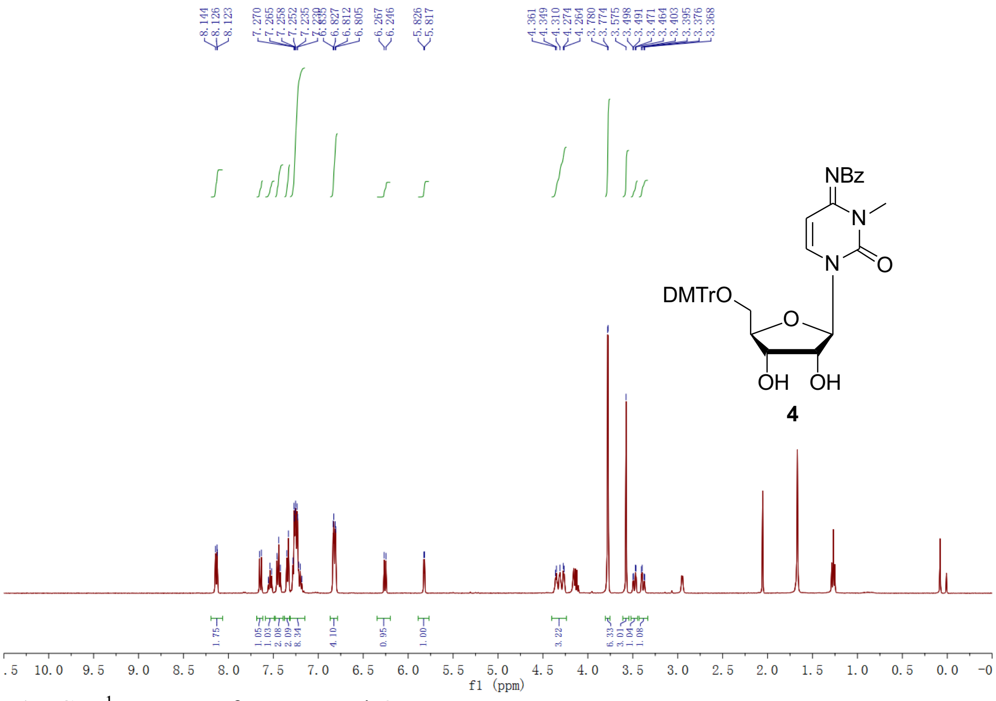

Fig. S7. [1]H NMR of compound **4**.

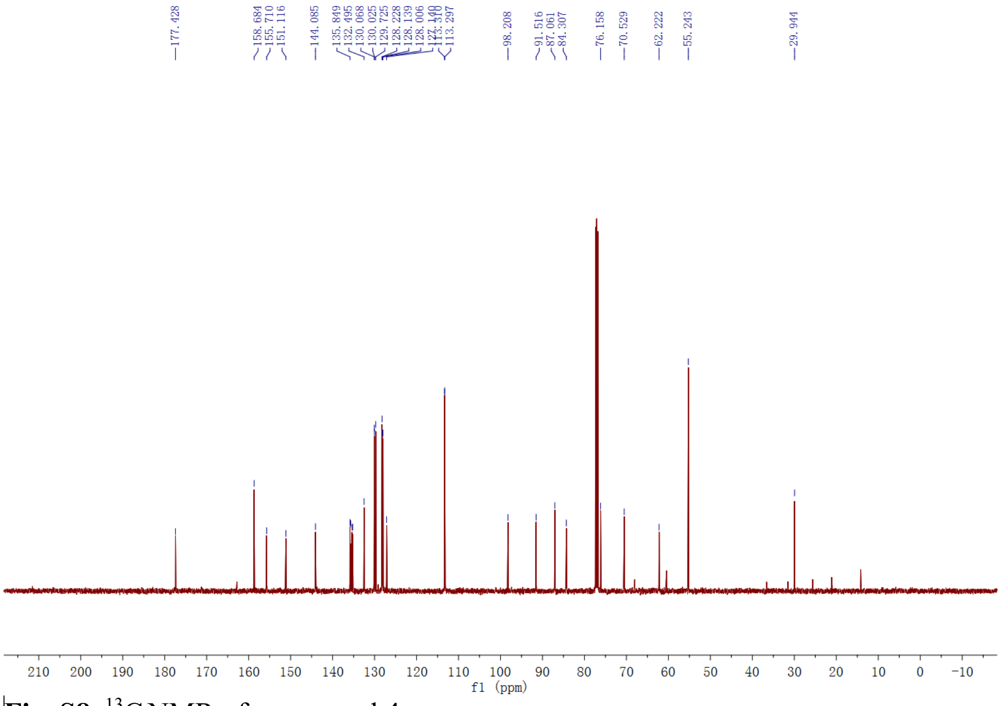

Fig. S8. [13]C NMR of compound **4**.

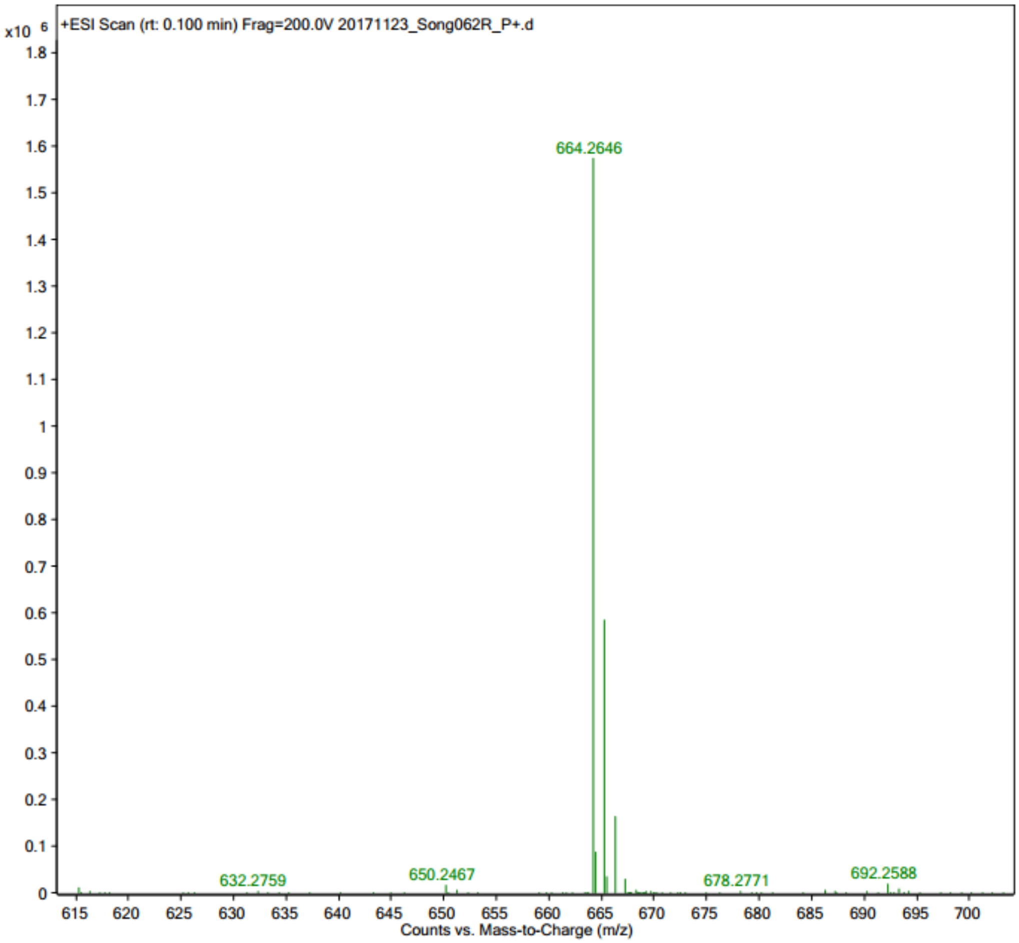


**Fig. S9.** Mass of compound **4**.

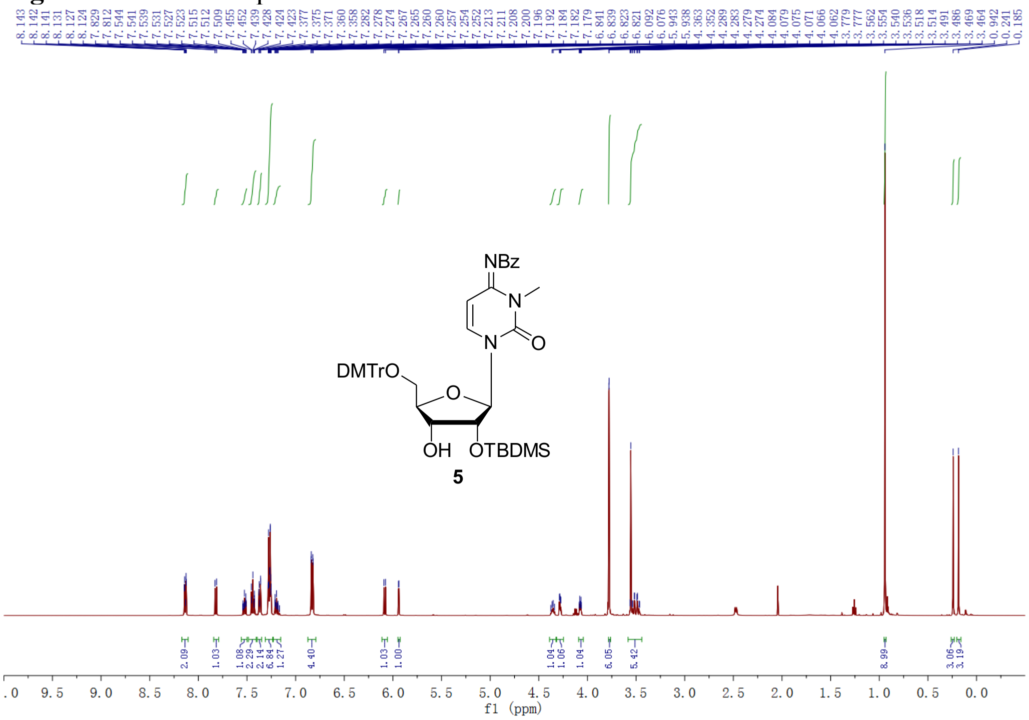

**Fig. S10.** $^{1}$H NMR of compound **5**.

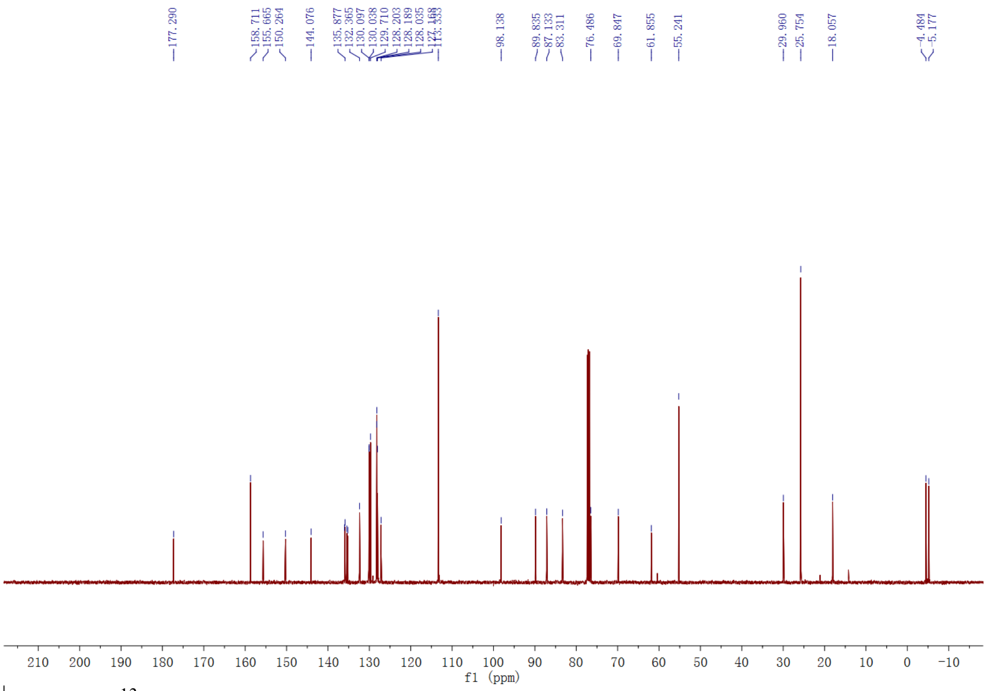


**Fig. S11.** $^{13}$C NMR of compound **5**.

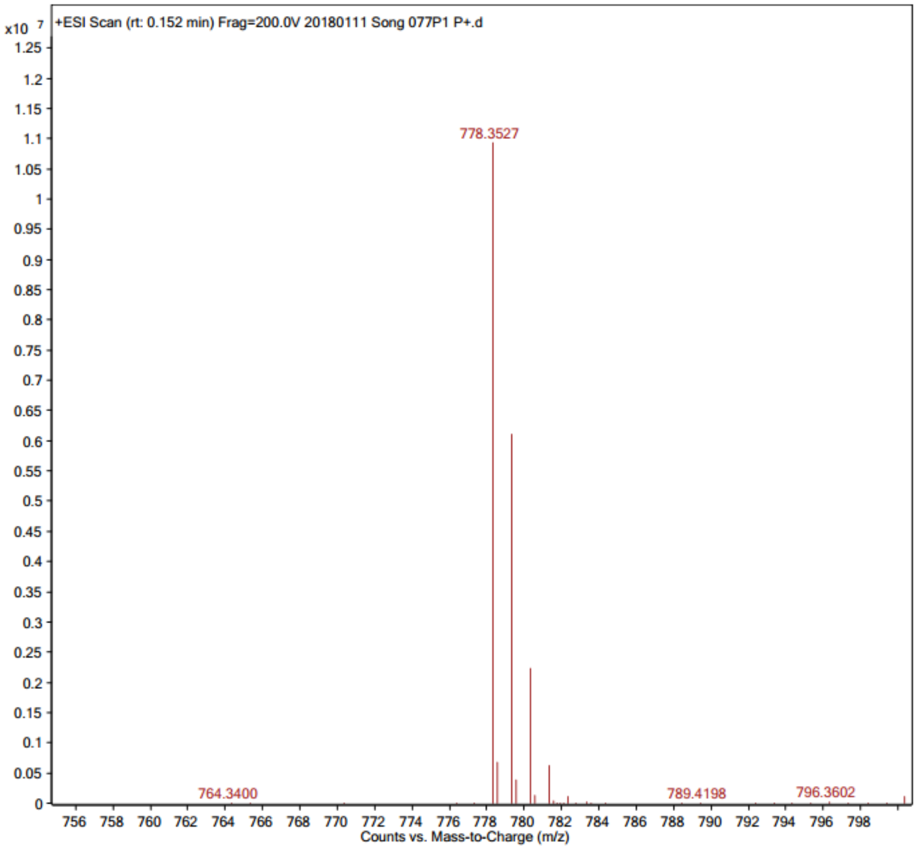

**Fig. S12.** Mass of compound **5**.

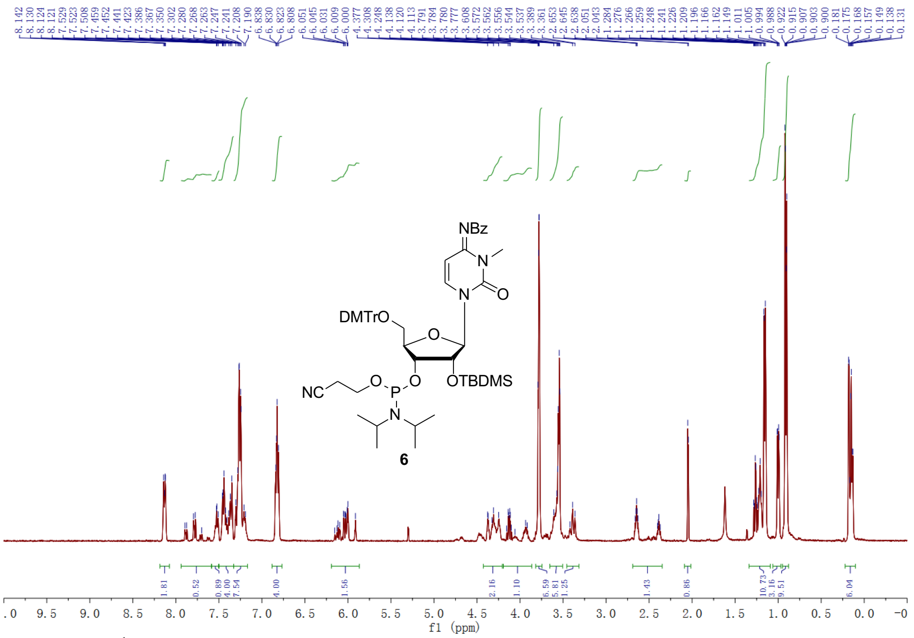


**Fig. S13.** $^{1}$H NMR of compound **6**.

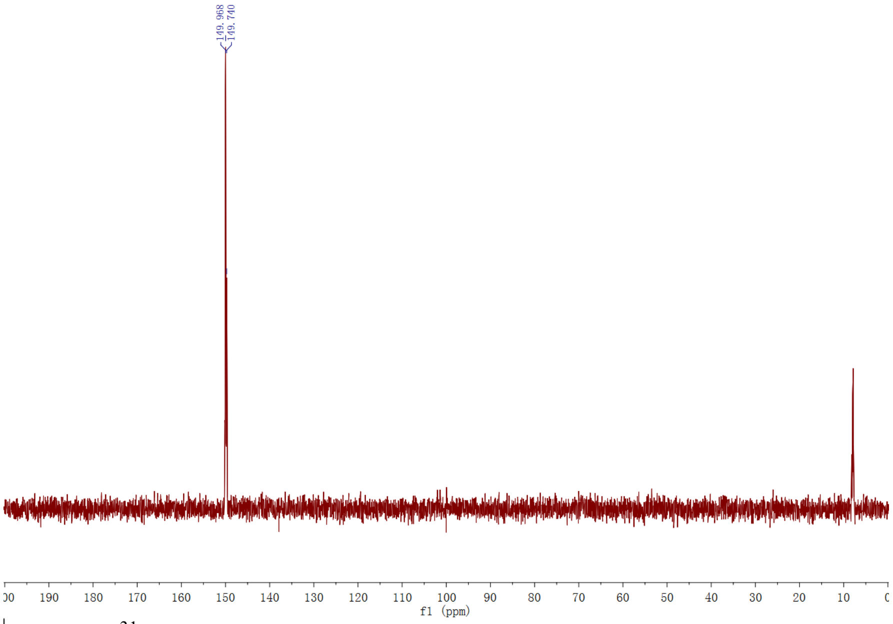


**Fig. S14.** $^{31}$P NMR of compound **6**.

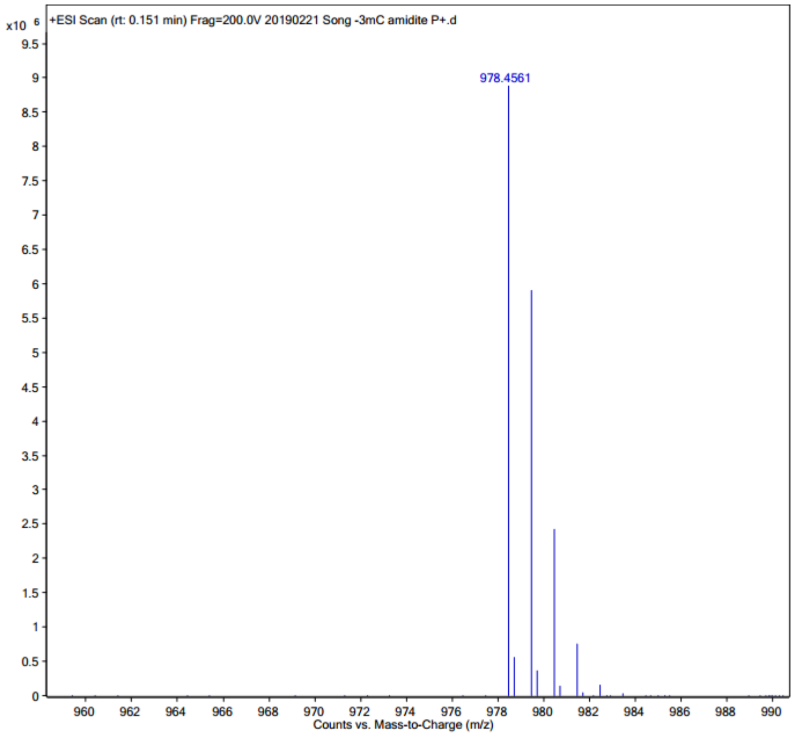


**Fig. S15.** Mass of compound **6**.

### Part II. Synthesis, HPLC and Characterization of RNA oligonucleotides

#### HPLC purification and analysis.

The oligonucleotides were purified by reverse phase HPLC using a Zorbax SB-C18 column at a flow rate of 1 mL/min. Buffer A was 20 mM Tris-HCl, pH 8.0; buffer B 1.25M NaCl in 20 mM Tris-HCl, pH 8.0. A linear gradient from 100% buffer A to 70% buffer B in 20 min was used to elute the oligos. The analysis was carried out by using the same type of analytical column with the same eluent gradient. All the modified-oligos were checked by high-resolution MS, as summarized in Table S1 and Fig. S19-33.

**Table S1.** RNA sequences containing $m^3C$.

| Entry | RNA Sequences | Measured (calc.) m/z |
|---|---|---|
| **ON1** | AAUGC$m^3$CGCACUG | $[M+H]^+$ = 3807.5 (3807.6) |
| **ON2** | GGACU$m^3$CCUGCAG | $[M+H]^+$ = 3823.6 (3823.6) |
| **ON3** | U$m^3$CGUACGA | $[M+H]^+$ = 2523.1 (2522.4) |
| **ON4** | GUA$m^3$CGUAC | $[M+H]^+$ = 2522.5 (2522.4) |
| **ON5** | CCGG$m^3$CGCCGG | $[M+H]^+$ = 3203.7 (3203.5) |
| **ON6** | CGCGAAUU$m^3$CGCG | $[M+H]^+$ = 3823.6 (3823.6) |

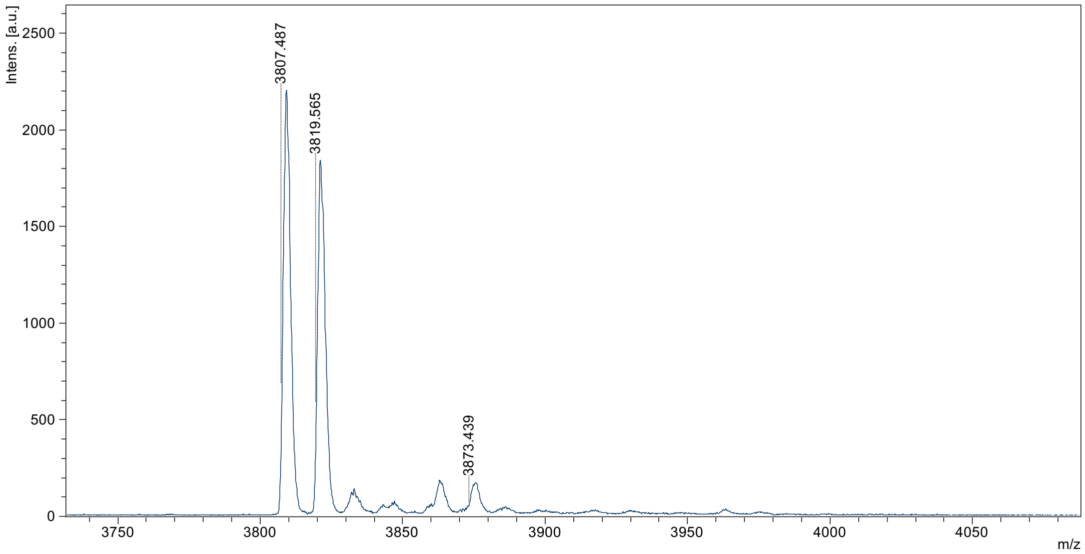


**Figure S19.** MALDI-TOF MS of **ON1** $m^3$C-12 mer (5’-AAUGC$m^3$CGCACUG-3’) $[M+H]^+$ = 3807.5 (calc. 3807.6).

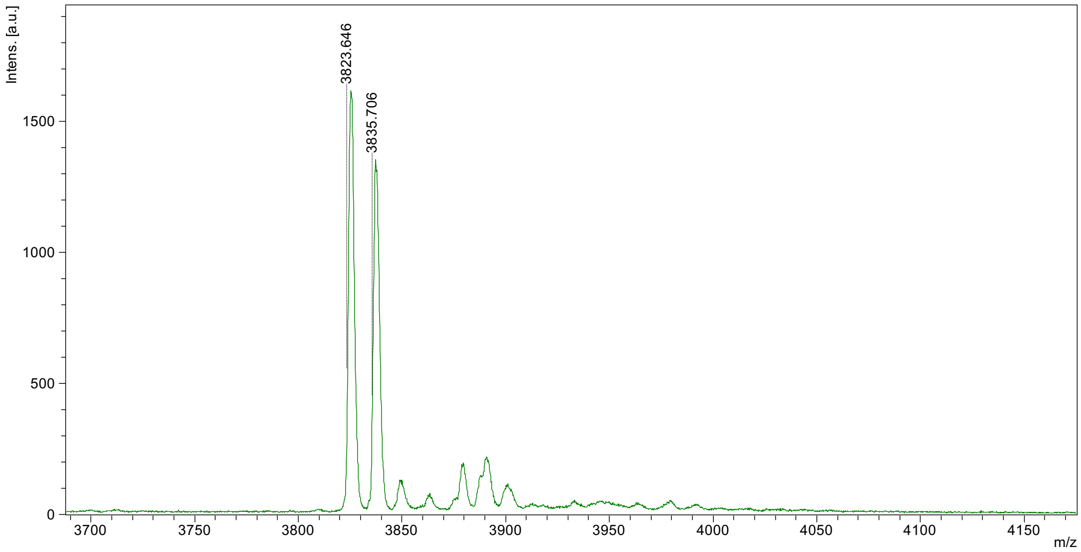


**Figure S20.** MALDI-TOF MS of **ON2** $m^3$C-12 mer (5’-GGACU$m^3$CCUGCAG-3’) $[M+H]^+$ = 3823.6 (calc. 3823.6).

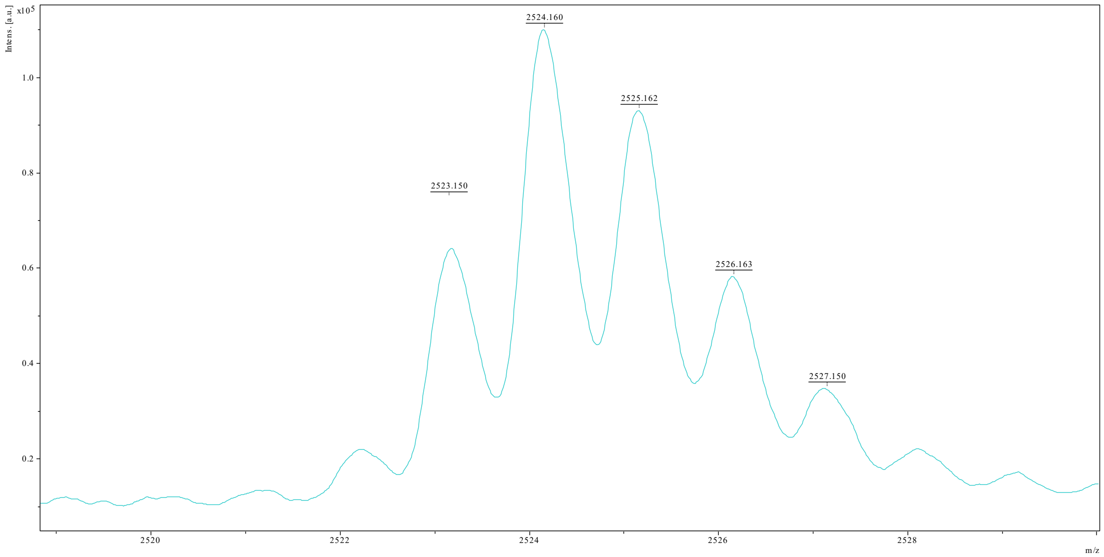


**Figure S21.** MALDI-TOF MS of **ON3** m$^3$C-8 mer (5'-Um$^3$CGUACGA-3') [M+H]$^+$ = 2523.1 (calc. 2522.4).

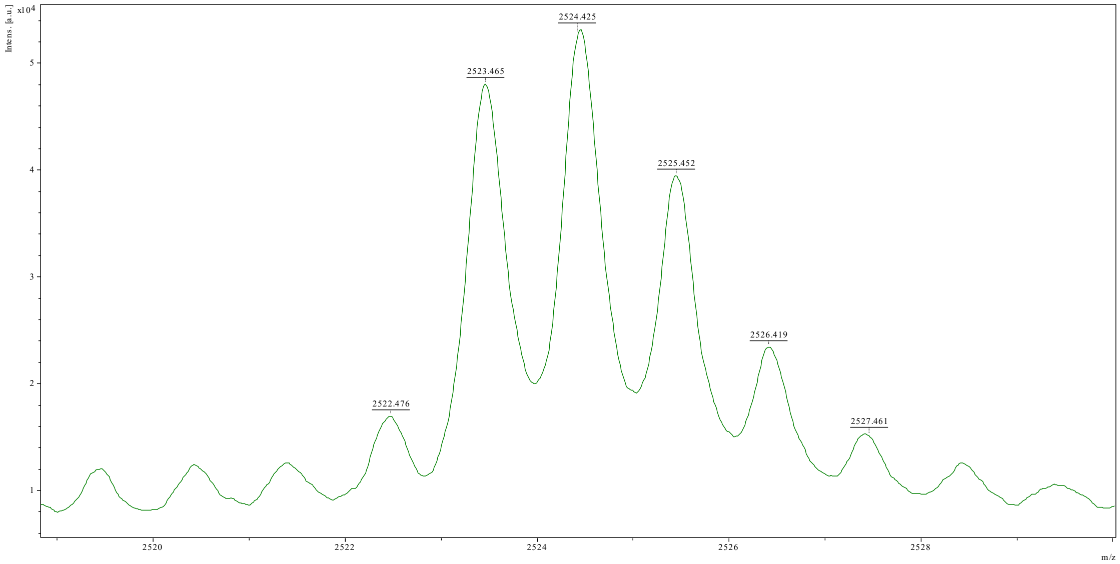


**Figure S21.** MALDI-TOF MS of **ON4** m$^3$C-8 mer (5'-GUAm$^3$CGUAC-3') [M+H]$^+$ = 2523.1 (calc. 2522.4).

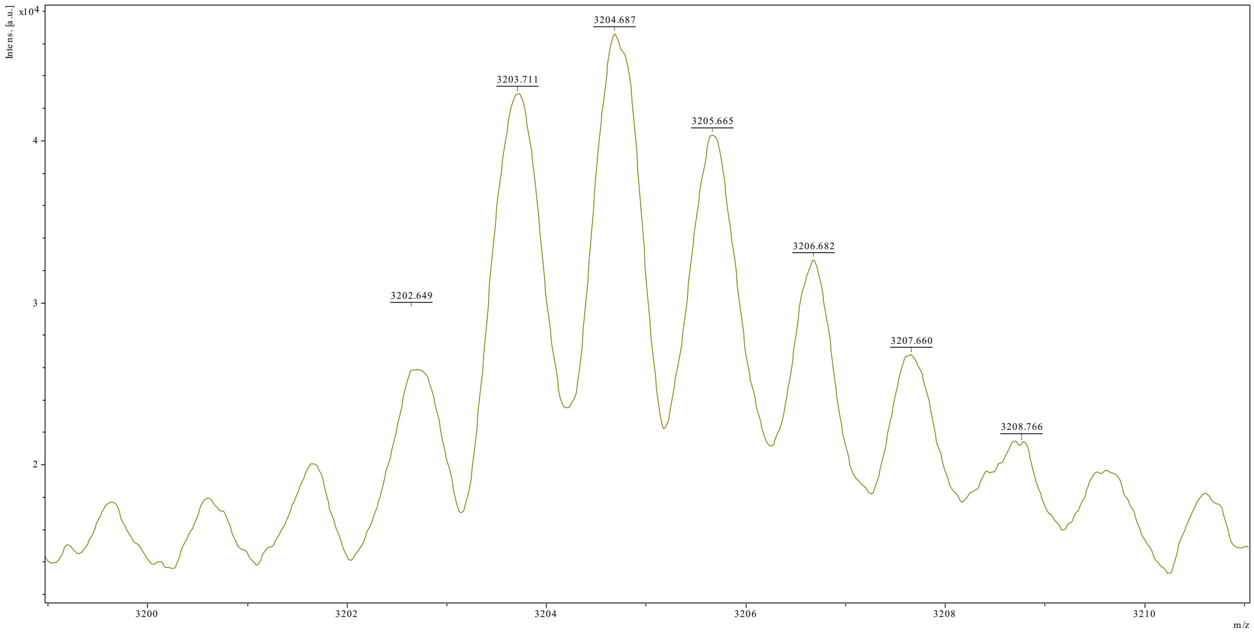


**Figure S21.** MALDI-TOF MS of **ON5** m$^3$C-10 mer (5'-CCGGm$^3$CGCCGG-3') $[M+H]^+$ = 3203.7 (calc. 3203.5).

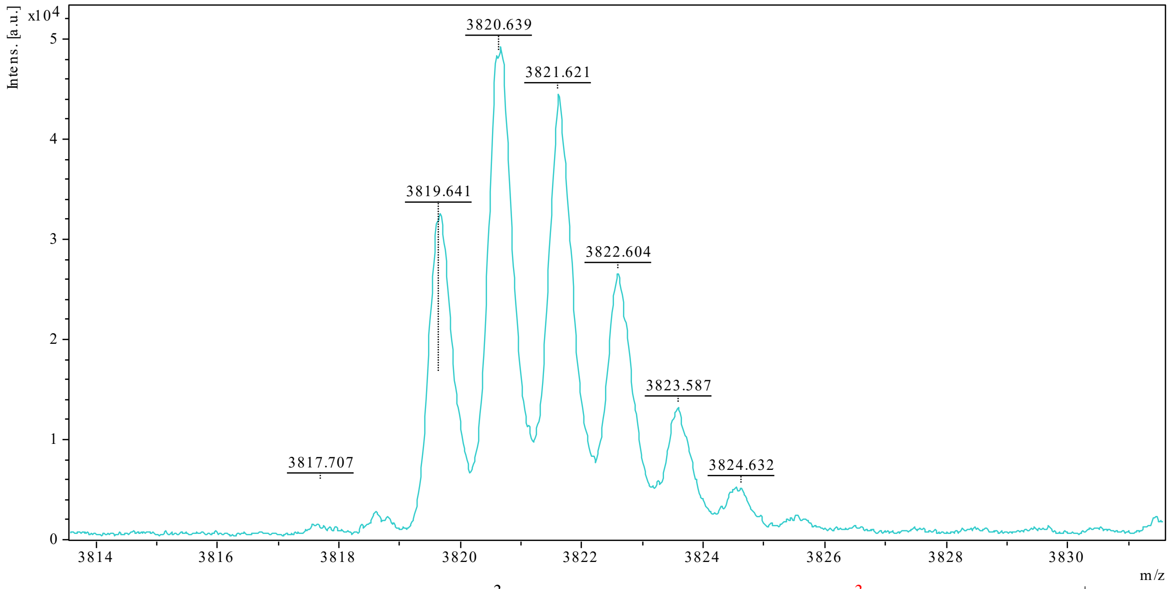


**Figure S22.** MALDI-TOF MS of **ON6** m$^3$C-12 mer (5'-CGCGAAUUm$^3$CGCG-3') $[M+H]^+$ = 3823.6 (calc. 3823.6).

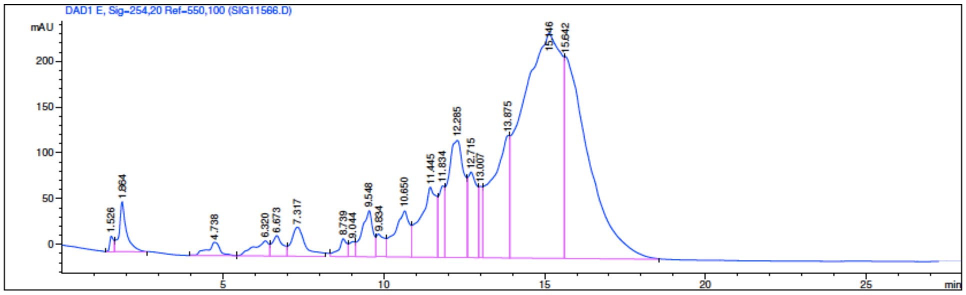


**Figure S27.** HPLC of **ON1** m$^3$C-12 mer (5'-AAUGCm$^3$CGCACUG-3')

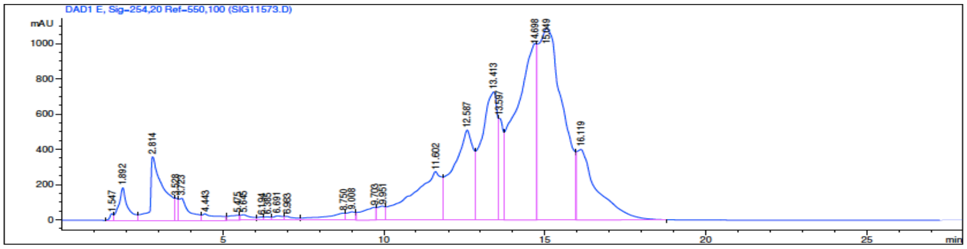


**Figure S28.** HPLC of **ON2** m$^3$C-12 mer (5'-GGACUm$^3$CCUGCAG-3')

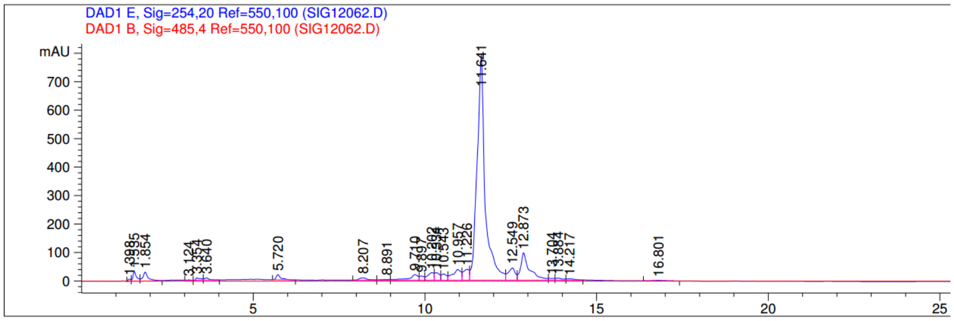


**Figure S29.** HPLC of **ON3** m$^3$C-8 mer (5'-Um$^3$CGUACGA-3')

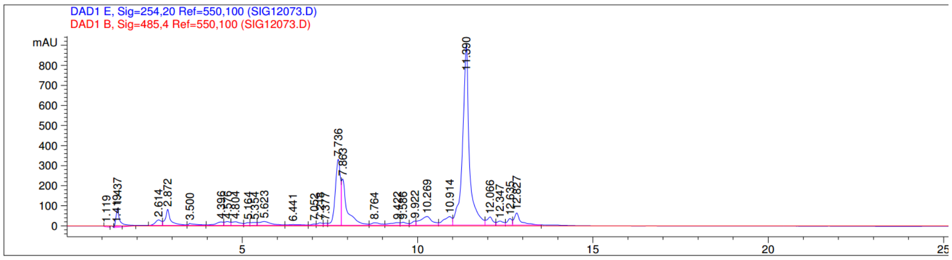


**Figure S30.** HPLC of **ON4** m$^3$C-8 mer (5'-GUAm$^3$CGUAC-3')

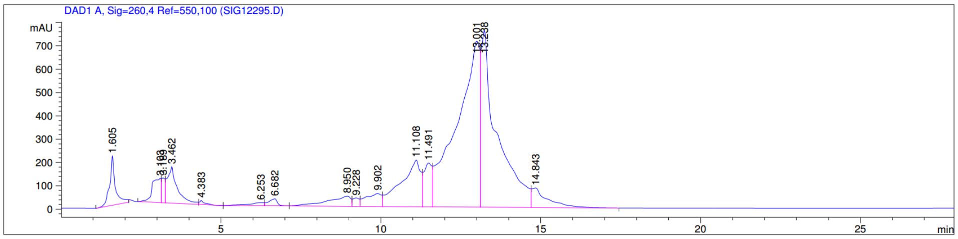


**Figure S31.** HPLC of **ON5** m$^3$C-10 mer (5’-CCGGm$^3$CGCCGG-3’)

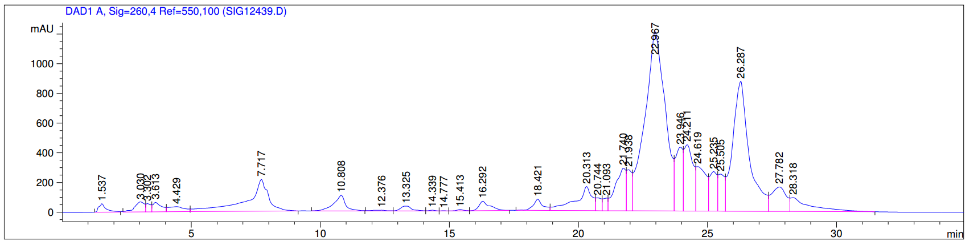


**Figure S32.** HPLC of **ON6** m$^3$C-12 mer (5’-CGCGAAUUm$^3$CGCG-3’)

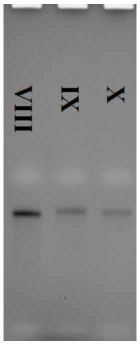


**Figure S33.** UV gel images of **VIII, IX and X-**22 mer oligonucleotieds containing one or two m$^3$C modifications..

### Part III. UV-melting temperature ($T_m$) study

Solutions of the duplex RNAs (1.5 µM) were prepared by dissolving the purified RNAs in sodium phosphate (10 mM, pH 7.0) buffer containing 100 mM NaCl. The solutions were heated to 95 °C for 5 min, then cooled down slowly to room temperature, and stored at 4 °C for 2 h before Tm measurement. Thermal denaturation was performed in a Cary 300 UV-Visible Spectrophotometer with a temperature controller. The temperature reported is the block temperature. Each denaturizing curves were acquired at 260 nm by heating and cooling from 5 to 80 °C for four times in a rate of 0.5 °C/min. All the melting curves were repeated for at least four times. The thermaldynamic parameter of each strand was obtained by fitting the melting curves in the Meltwin software.

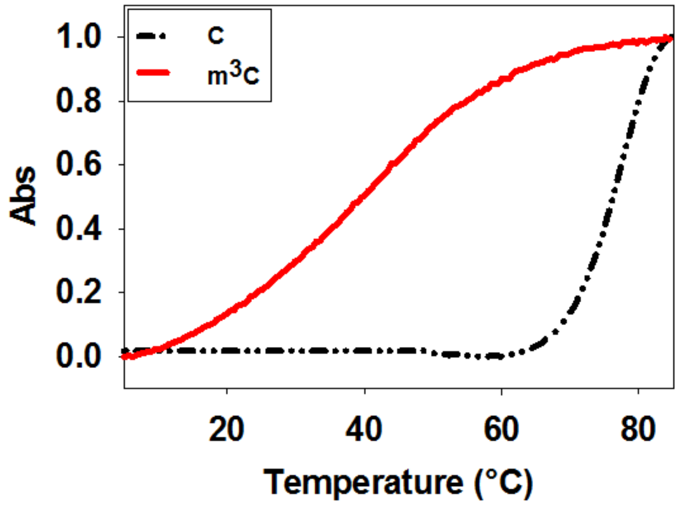


**Figure S34.** Normalized UV-melting curves of self-complementary RNA duplex. Native 10-mer sequence 5′-CCGGCGCCGG-3′ strand (in black line) and $m^3C$ modified 5′-CCGG$m^3C$GCCGG-3′ strand (in red line).

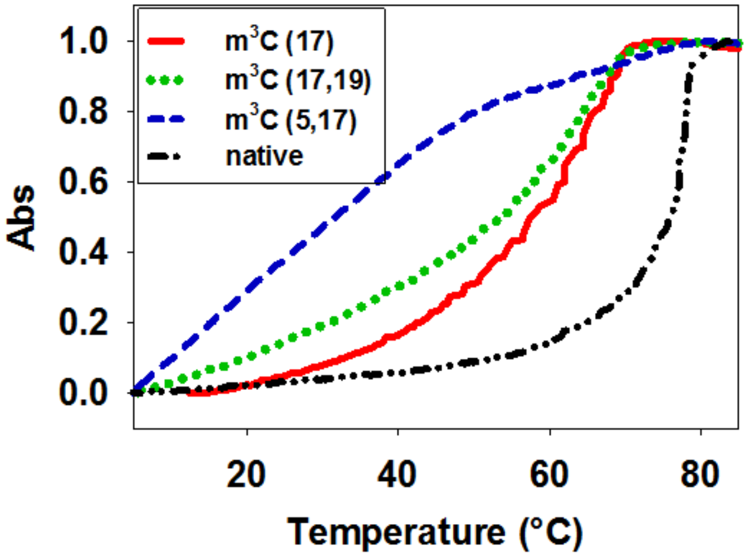


**Figure S35.** Normalized UV-melting curves of the 22-mer 5′-UGAGCUAGUAGGUUGUCUCGUU-3′ with one or two $m^3C$ modifications in various position.

**Part IV. UV gel images of standing-start primer extension reactions.**

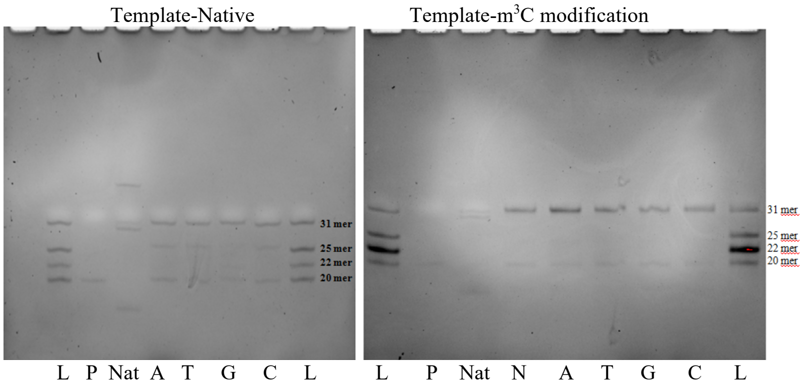


**Figure S36.** UV gel images of standing-start primer extension reactions for AMV RT indicated using m$^3$C containing RNA template and the corresponding natural template. Lanes: L, ladders; P, primer; Nat, natural template with all four dNTPs; A, T, G, and C, reactions in the presence of the respective dNTP; N, reactions in the presence of all four dNTPs.

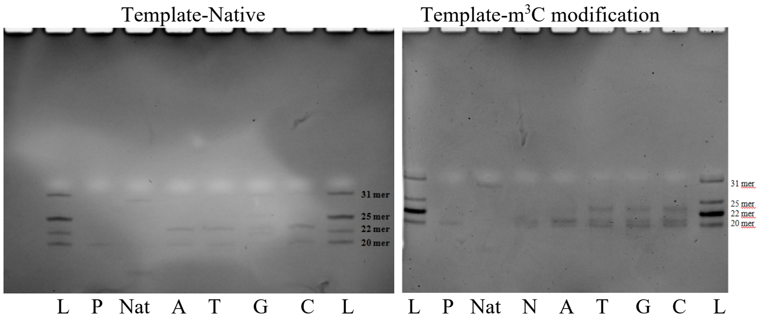


**Figure S37.** UV gel images of standing-start primer extension reactions for HIV-1 RT indicated using m$^3$C containing RNA template and the corresponding natural template. Lanes: L, ladders; P, primer; Nat, natural template with all four dNTPs; A, T, G, and C, reactions in the presence of the respective dNTP; N, reactions in the presence of all four dNTPs.

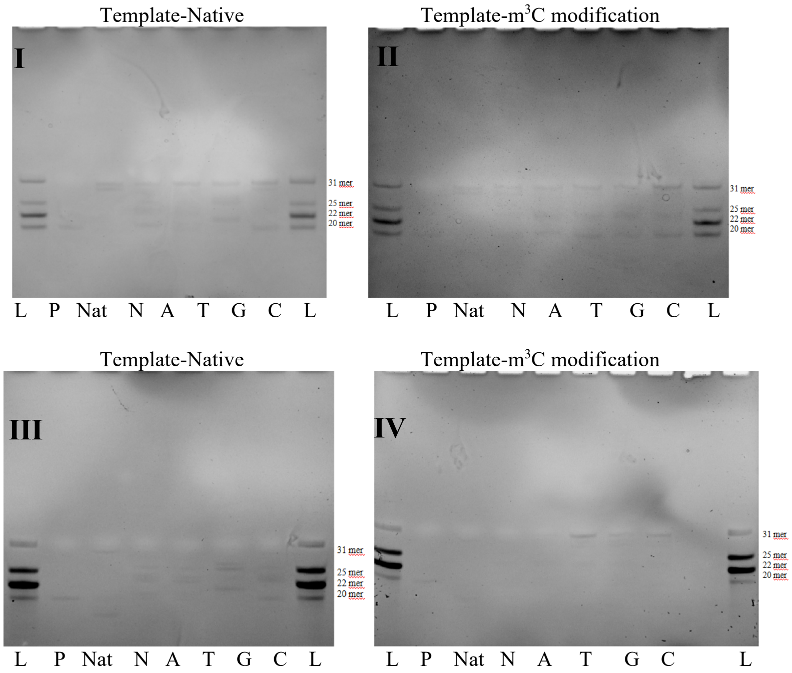


**Figure S38.** UV gel images of standing-start primer extension reactions for MMLV RT (**I** and **II**) and MultiScribe$^{TM}$ RT (**III** and **IV**) indicated using m$^3$C containing RNA templates and the corresponding natural templates. Lanes: L, ladders; P, primer; Nat, natural template with all four dNTPs; A, T, G, and C, reactions in the presence of the respective dNTP; N, reactions in the presence of all four dNTPs.
